# Temperature stress disrupts reciprocal adaptation in a microbial predator-prey system

**DOI:** 10.1101/2025.11.21.689813

**Authors:** Karissa Plum, Rebecca Zufall

## Abstract

Antagonist interactions, such as predator-prey interactions, are widespread in nature and drive both ecological and evolutionary outcomes. Coevolutionary outcomes of antagonistic interactions have been shown to be influenced by environmental conditions, yet the role of abiotic stress in modifying these outcomes remains insufficiently understood. Here we explored how the addition of temperature stress altered evolutionary trajectories of traits of both species in the *Pseudomonas fluorescens – Tetrahymena pyriformis* (bacteria-ciliate) predator prey system. We found that temperature stress impeded the evolution of traits important for antagonistic interactions in both species. Prey defense levels as well as predators’ ability to eat prey were limited under temperature stress. We also found that the addition of temperature stress altered growth rate evolution in evolving populations of both species. Taken together, our results show that temperature stress not only alters the evolutionary trajectories of both predator and prey traits but also hinders their coevolution. These findings suggest that environmental stressors may weaken reciprocal coevolution which could have important consequences for the stability and persistence of ecological communities.

## Introduction

Antagonistic interactions, such as predator-prey or host-parasite interactions, are ubiquitous in nature and have been shown to drive both ecological processes (e.g. population dynamics; Fussmann et al., 2000; Getz & Owen-Smith, 2011; Kingsland, 2015; Paine, 1966) and evolutionary outcomes (Abrams, 2000; Burak et al., 2018; Thompson, 2005). In predator-prey systems, coevolution often occurs as a result of reciprocal selection on interacting species.

Antagonistic coevolution in these systems occurs when prey evolve defenses against predators and predators evolve countermeasures to overcome those defenses. This can result in arms race dynamics where both species evolve increasing values of traits involved in defense/countermeasures over time (Buckling & Rainey, 2002; Dawkins & Krebs, 1997; Harrison et al., 2013; Thompson, 2005) or fluctuating selection dynamics where trait values oscillate (Abrams & Matsuda, 1996; Borghans et al., 2004; Dieckmann et al., 1995; Nuismer & Thompson, 2006; Seger, 1988).

The ecological and evolutionary outcomes of antagonistic predator-prey interactions often depend on environmental conditions. In particular, abiotic stress has the potential to alter, e.g., interaction strength, behavior, and phenotypes of predators and prey (Tylianakis et al., 2008). Increased temperatures have been shown to affect a variety of traits associated with predator-prey interactions, including sensory performance, handling time, attack/killing rates, and non-consumptive behaviors (Draper & Weissburg, 2019; Janssens et al., 2015; Öhlund et al., 2015; Robertson & Hammill, 2021; Walker et al., 2020). Further, the addition of abiotic stress into predator-prey interactions has been shown to change evolutionary outcomes by altering the direction, strength, or pace of both coevolutionary and abiotic adaptation (Govaert et al., 2019; Northfield & Ives, 2013; Theodosiou et al., 2019). For example, a study using a C*haoborus americanus* larvae - *Daphnia pulex* predator-prey system found that the addition of predation altered thermal adaption in prey, possibly due to selection for higher growth rates in prey populations experiencing predation stress (Tseng & O’Connor, 2015). Theoretical models predict that increased prey productivity enhances the evolution of defensive traits and accelerates arms race dynamics (Hochberg & Baalen, 1998), and experimental studies in a bacteria-ciliate predator-prey system have shown that enriched resources can promote prey defense and weaken predator control (Friman et al., 2008). Additionally, in a bacteria-phage system, fluctuating nutrient availability was shown to constrain coevolution when environmental fluctuations occurred faster than selective sweeps could occur (Harrison et al., 2013). For a full understanding of predator-prey coevolution, it is thus necessary to examine the effects of abiotic factors on these antagonistic interactions.

Microbial systems make excellent models to study the effects of abiotic stress on coevolution in predator-prey systems for a variety of reasons. First, microbes are important members of all natural communities, playing pivotal roles in nutrient cycling, animal and plant health, and global food webs (Cavicchioli et al., 2019; Gougoulias et al., 2014; Redford, 2023). Smaller microbes, such as bacteria and algae, serve important ecological roles such as plant-soil feedbacks, carbon sinks, and food sources for larger organisms (Gupta et al., 2016; Jassey et al., 2022; Pandey, 2023). Larger microbes, such as ciliates and flagellates, function as important links in various food webs, feeding on smaller microbes and serving as prey to zooplankton and small vertebrates (Finlay & Esteban, 1998; Jack & Gilbert, 1997; Porter et al., 1979; Sherr & Sherr, 2002; Vander Zanden & Vadeboncoeur, 2002). Second, predator-prey interactions among microbes exhibit many similarities to those in animals, including foraging/prey capture behavior and predator evasion behavior (Schmitz, 2017). This means that results from microbial studies are likely to be applicable to multicellular systems as well. Finally, microbial predator-prey systems are experimentally tractable in lab settings. Experimental evolution, wherein populations are allowed to evolve under highly controlled laboratory conditions, is becoming more frequently used to study microbial predator-prey interactions (Brockhurst & Koskella, 2013; McDonald, 2019). Many microbes are well suited for this type of experiment due to short generation times, ease of culture in lab, and the ability to store intermediate populations (e.g., by freezing; (Bennett & Hughes, 2009; Kawecki et al., 2012; Plum et al., 2022).

A powerful system previously developed for use in microbial predator-prey experimental evolution is the *Tetrahymena* – *Pseudomonas* system. *Tetrahymena* are bactivorous filter-feeding ciliated microbial eukaryotes found globally in freshwater ecosystems (Doerder, 2019; Doerder & Brunk, 2012; Eisenmann et al., 1998; Simon et al., 2008; Thurman et al., 2010). *Pseudomonas* are flagellated bacteria, commonly found in both soil and water, as well as associated with animal and plant hosts (Mei et al., 2024; Rhodes, 1959; Shalev et al., 2022; Silby et al., 2011; Zhou et al., 2021). In response to predation, *Pseudomonas* have been shown to evolve the ability to form biofilms or produce toxic compounds (Hoque et al., 2023; Jousset et al., 2006, 2009; Matz & Kjelleberg, 2005; Seiler et al., 2017; Weitere et al., 2005). In the presence of prey, *Tetrahymena* evolve changes in morphology and feeding behavior, such as swimming patterns (Cairns et al., 2020). Studies of eco-evolutionary dynamics in this system have examined how adaptation alters population dynamics (Friman et al., 2014; Hiltunen et al., 2015, 2018; Hiltunen & Becks, 2014; Scheuerl et al., 2019). For example, the presence of antibiotics hinders adaptation to predators, altering population cycles (Hiltunen et al., 2018). Studies that address coevolution in this system often use coevolved predators and/or prey lines generated in earlier experiments and test how evolutionary history effects community dynamics including population cycles and gene expression patterns (Hiltunen & Becks, 2014; Hogle et al., 2023; Scheuerl et al., 2019). To our knowledge, only one study reports the evolutionary effects of abiotic stress on both the predator and prey in this system. That study found that defensiveness of prey evolved more slowly in the presence of fluctuating nutrients and salinity. However, the abiotic environment did not affect predator evolution; predators evolved counterdefenses regardless of the environmental stress (Hiltunen et al., 2015). These studies demonstrate both the utility of this model system and the need to further address the role of abiotic conditions on both partners in antagonistic coevolution.

To better understand the evolutionary consequences of abiotic stress on predator-prey coevolution, we used experimental evolution to study the effects of thermal stress on trait evolution in coevolving populations of *T. pyriformis* and *P. fluorescens*, which initially have similar thermal optima. For both species, we measured defense or predation efficiency as well as growth rates, as traits that are likely to evolve under temperature stress and predator-prey interactions. Consistent with our predictions, we found that temperature stress hindered the evolution of traits important for species interactions in both species. In addition, contrary to most previous microbial evolution experiments, we did not find consistent increases in growth rates in response to either increased temperature or adaptation to laboratory conditions in either species. Overall, our results indicate that reciprocal adaptation in this system is hindered by temperature stress.

## Methods

### Study System

A derivative of the bacterial strain *P. fluorescens* SBW25 (Rainey & Travisano, 1998) was the prey species used in this experiment and the ciliated protist *T. pyriformis* strain GL-C (Tetrahymena Stock Center SD00707) was the predator. Bacteria were grown in Lauria Broth (LB) medium (10 g/L tryptone, 5 g/L yeast extract, and 5 g/L NaCl) and stored at -80°C. When not co-cultured with *P. fluorescens*, ciliates were grown in standard nutrient-rich *Tetrahymena* medium, SSP (Cassidy-Hanley, 2012; Gorovsky et al., 1975) and were stored under liquid nitrogen.

### Experimental Evolution

To minimize genetic variability in starting populations, both predator and prey populations were started from fully clonal populations. These were generated by picking a single colony (for bacteria) or cell (for ciliates) and growing in liquid medium for 24 or 48 hours respectively. These cultures were then used to start experimental evolution populations. Subsamples from these cultures were also frozen to use as the ancestor for trait assays.

Four replicate populations of *P. fluorescens* and *T. pyriformis* were evolved with one species in a culture, which we call “alone,” or with both species, termed “together,” at 26°C, close to their thermal optimum, or 30°C, closer to their thermal maximum (Figure 1A), for a total of 24 populations. All populations were evolved in 250 ml flasks with 50 ml of medium. Alone populations of *T. pyriformis* were cultured using 5% SSP; alone populations of bacteria and together populations were grown on 5% LB. All populations were started with the same densities for each organism: 1000 cells/ml for *T. pyriformis* and ∼3,200,000 cells/ml for *P. fluorescens*. Ten percent of each flask was moved to fresh medium twice a week. This subculture regime resulted in all populations reaching stationary phase prior to transfer. Populations evolved for approximately 6 months, ∼508-768 prey generations or ∼209-829 predator generations (Supplementary Table S1). Generation ranges were estimated assuming that no cell division happens during stationary phase. Since this assumption is likely unrealistic, generation ranges should be taken as rough approximations.

**Figure 1.**
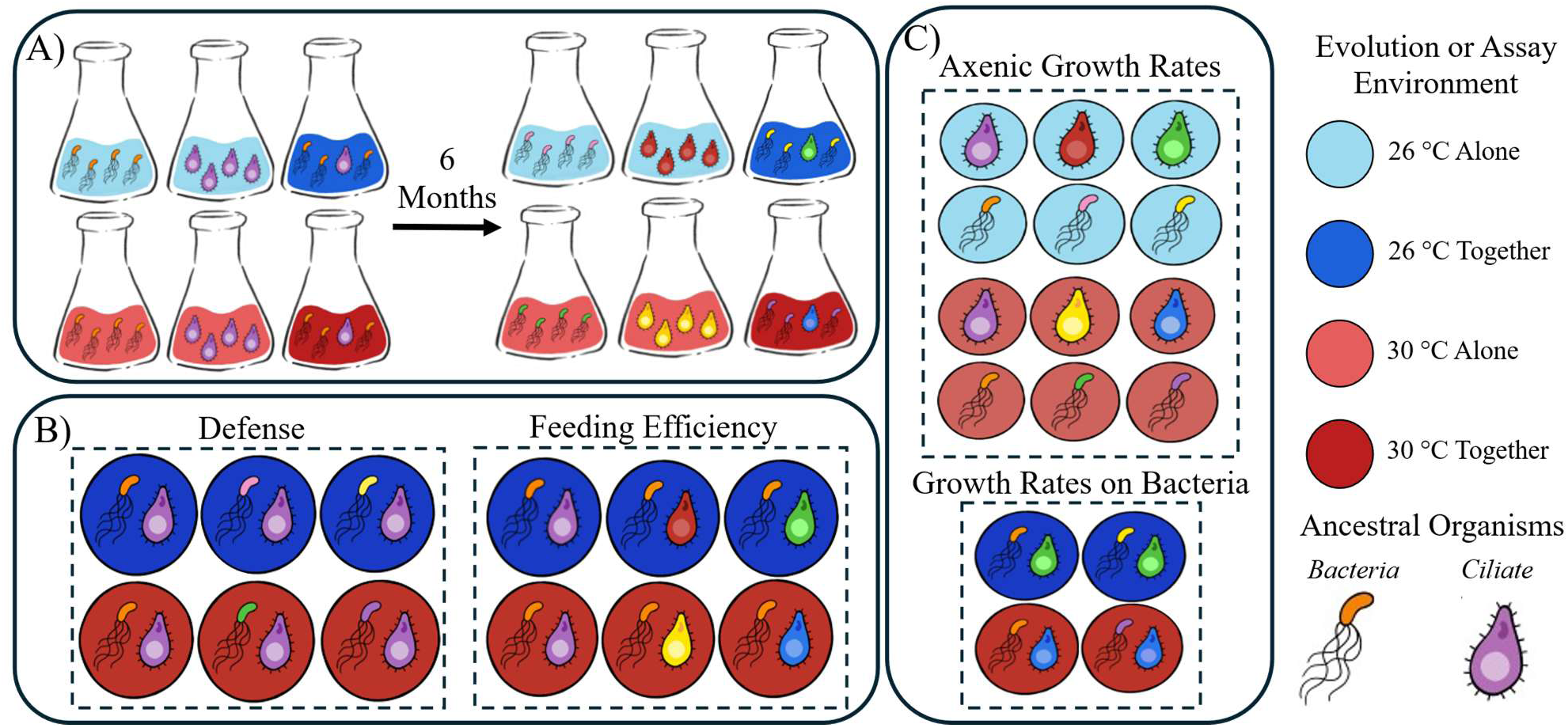
Experimental Design. (A) All experimental populations were started from clonal isolates of bacteria and ciliates. Ancestral populations of bacteria and ciliates are represented with orange and purple organisms respectively. Evolutionary histories of each organism is indicated by different colored individuals after 6 months of evolution. Populations evolved at 26°C (blue) or 30°C (red) either alone (faded) or together (darker shades) for 6 months. (B) Defensiveness and feeding efficiency of evolved populations were measured at all 6 timepoints (defense) or the final timepoint of the experiment (feeding efficiency). Ancestor partners were used to quantify traits of evolved populations, and trait values were quantified relative to the ancestor populations (see methods). C) Growth rates for both species were quantified in axenic medium. Evolved populations were assayed at their evolution temperature, and ancestors were assayed at both temperatures. Growth rates on bacterial prey were also qualified for only the coevolved predator lines. Growth rates for theses predators were assayed using both ancestral and coevolved bacteria.

Samples from all populations were saved monthly. Bacteria were saved by making 20% glycerol stocks and storing them at -80°C. Ciliates were frozen in liquid nitrogen using the protocol from Cassidy-Hanley et al. (1995). Ciliate samples from together evolved populations were first treated with antibiotics to eliminate bacteria before freezing. Ciliates do not survive freezing in glycerol, so the together bacteria populations did not need further treatment prior to freezing.

### Defensiveness assay

Defensiveness of bacteria was quantified using an assay that measures how well ancestral predators are able to eat and use prey evolved under different conditions to increase their population sizes (Hiltunen & Becks, 2014). Following the experimental evolution, prey from all evolution conditions as well as the ancestor were thawed; 100ul was inoculated into fresh 5% LB medium; and cultures were grown shaking at room temperature (∼21°C). After 24 hours of growth, OD was measured, and each culture was diluted to OD ∼0.2A to ensure similar starting densities for all populations. Once target ODs were reached, 100ul of the diluted sample was used to inoculate 2ml of fresh 5% LB medium in a 24 well plate, with 4 replicates per population. To start the assay, 2100 individuals from the ancestral predator stock were added to each well. Before adding the predators, the high nutrient SSP medium was removed by centrifugation to ensure that population growth of predators is due to consuming bacteria and not due to carry over of the ciliate medium (Figure 1B). After 48 hours, predator numbers in each well were counted using a dissecting microscope. Relative defensiveness was calculated as 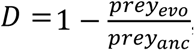, where *prey_evo_* is the predator density of cultures fed evolved prey, and *prey_anc_* is the predator density when fed evolved prey (Supplementary Table S2). Values close to zero indicate little to no change in the defense level of the evolved prey compared to the ancestor. Large, positive values indicate a high level of evolved defense. Defense was measured from samples saved at every timepoint in the experiment.

### Feeding Efficiency assay

Feeding efficiency of the ciliates was measured similarly to defensiveness to quantify how well evolved predators consume and metabolize ancestral prey to increase their population sizes. Evolved and ancestral predator lines were thawed from liquid nitrogen (Cassidy-Hanley, 2012) and grown axenically in SSP at their evolution temperatures. Ancestor lines were kept at 26°C, the ancestral thermal optimum. Once populations were reestablished, 250ul of each population was used to inoculate 5.5 ml of fresh SSP medium and were left to grow at their evolution temperature until carrying capacity was reached (2-3 days). One day before the assay plate was set up, ancestral bacteria stock was grown as described above (Defensiveness Assay). Assay plates were set up by adding 100ul of ancestral prey culture at OD 0.2 and 2100 predators (4 replicates per evolved predator population; Figure 1B). The number of ciliates in each culture were counted after 48 hours. Relative feeding efficiency was calculated as 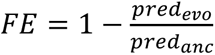, where *prey_evo_* is the density of evolved predators that were fed ancestral prey, and *prey_anc_* is the density of ancestral predators fed ancestral prey (Supplementary Table S3).

### Growth Rate Assay on Axenic Medium

Growth curves for both species were generated by inoculating ∼1000 cells of *P. fluorescens* or ∼500 cells *T. pyriformis* into a well on a 96-well plate containing LB or SSP, respectively, then measuring the optical density (OD) at 650 nm in a microplate reader at their evolution temperature every 5 min for 72 hours. Populations from both species evolved at the same temperature from a single timepoint were assayed on the same plate and there were 5 replicates measured per population. Growth curves for ancestral populations of both species were measured as an internal control on every plate. Because not all ciliate timepoints survived freezing, only some timepoints were measured for *T. pyriformis*. All timepoints were assayed for *P. fluorescens*, except for 26°C evolved populations at the second time point due to plate reader errors (Figure 1C).

Growth curves were analyzed using the R package growthrates (Petzoldt, 2025). Multiple models were fit to the growth curves and compared using AIC. The logistic model (Nelder, 1961) provided the best overall fit for all the growth curves of both species. Growth rates, in units of divisions/hour, were then extracted from the fit curves (Supplementary Table S4 & S6).

### Growth Rate of Predators on Bacteria

Growth rates of coevolved ciliates feeding on bacteria were also measured to test whether food type affects growth rate (Supplementary Table 8). Coevolved and ancestral populations of ciliates were fed ancestral bacteria or the bacteria with which they had coevolved. Assay plates were set up as described above (Feeding efficiency assay) by adding 100ul of prey at OD 0.2 and 2100 predators (4 replicates per predator population). Plates were incubated at the predator’s evolution temperature and ciliates were counted after 48 hours.

Doubling time was calculated using: 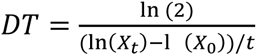 (Monod, 1949) where, *X_t_* = Cell density after 48 hours, *X*_0_ = Starting cell density, *t* = time(hrs).

### Data Analysis

#### Defense and Feeding Efficiency at final timepoint

We analyzed the effects of temperature (T), whether evolution was alone or together (I), and their interaction (T×I) on the evolution of bacterial defense (D) and ciliate feeding efficiency (FE) at the final timepoint of evolution. Assumptions of normality and homoscedasticity were checked using model residuals.

We fit a generalized linear mixed model (GLMM) using the R package glmmTMB to the defense data, with replicate population as a random effect and gaussian family and identity link, allowing variance to differ among evolution conditions using dispformula (Brooks et al., 2017). Model adequacy was assessed with DHARMa simulated residuals (Hartig et al., 2024). Fixed effects of evolution conditions were obtained using Type-III Wald χ² using the ANOVA function in the R package car (Fox et al., 2024).

For feeding efficiency, we fit a linear mixed effect model with replicate population as a random effect using the lmer function of the R package lme4 (Bates et al 2015). Effects of evolution conditions were obtained using Type-III ANOVA using Kenward-Roger degrees of freedom (Kuznetsova et al., 2017) and car (Fox et al., 2024).

Models with various combinations of (T) and (I) were fit and compared using AICc. The full interactive model was the best fit for both traits:

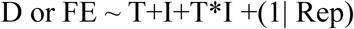

We estimated marginal trait means using r package emmeans (Lenth et al., 2025). Pairwise comparison of marginal means using Bonferroni correction for multiple comparisons was done as a post hoc test for difference in means trait values. T-tests were used to test if change in trait means of populations evolved in a similar condition differed from zero.

#### Defense evolution trajectory

The evolutionary trajectory of defensiveness over the course of the experiment was analyzed using a generalized additive mixed model (GAMM). The model was fit with separate smooth terms over time for each evolution condition using s(Days, by = group), including random effects due to biological replication using the gamm function of the R package mgcv (Wood, 2017). Initial data exploration of model fit residuals indicated heteroskedasticity of residuals, so we allowed variance to differ by evolution condition using varIdent. Models with various combinations of (T) and (I) were fit and compared using AICc. The full interactive model was the best fit:

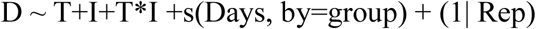

In order to test if there was a difference in the defense trait trajectories of populations evolved with predators at different temperatures, slopes of trait trajectories were obtained by calculating first derivatives of all smooth terms using the derivatives function in the R package gratia (Simpson, 2024). 95% confidence intervals around first derivatives were then assessed to identify time periods during which confidence intervals did not include zero (Simpson, 2018). This indicated periods of time where growth rates are changing.

#### Growth rate

Change in growth rate (Δr) was calculated for each replicate population as the population growth rate minus the average growth rate of the ancestor assayed at that time point (Supplementary Table S5 & S7). Change in growth rates for each species were analyzed individually. Data were analyzed similar to defense trait over time, using a regular GAMM (unweighted) on the *T. pyriformis* data and using a weighted GAM to account for heteroskedasticity in the *P. fluorescens* data. We included separate smooth terms over time per evolution condition and random effects due to biological replication in both models. Models with various combinations of (T) and (I) were fit and then compared using AICc. Model fits were evaluated by using the gam.check function of the mgcv package R (Wood, 2017). The full interactive model was the best fit for *P. fluorescens* trajectories while a model only including species interactions fit best for *T. pyriformis* trajectories:

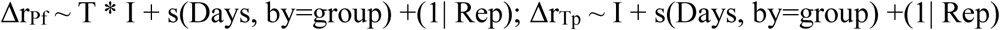

Periods of time where change in growth rate was significantly increasing or decreasing were identified as with defense trait trajectories.

#### Predator growth rate on bacteria

Change in growth rate (ΔDT) was calculated for each replicate population as the population doubling time minus the average doubling time of the ancestor assayed at that time point (Supplementary Table S9). We analyzed the effects of temperature (T), bacterial prey identity (B) i.e. ancestral or coevolved, and their interaction (T×B) on the evolution of doubling times of coevolved ciliate predators when fed bacterial prey using linear mixed effect models with random effects due to biological replication using the lmer function of the R package lme4. Models with various combinations of (T) and (B) were fit and then compared using AICc. Model fits were evaluated by using the gam.check function of the mgcv package R (Wood, 2017). The full interactive model was the best fit:

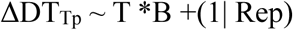

Effects of evolution condition were obtained using Type III ANOVA using Kenward-Roger degrees of freedom (Kuznetsova et al., 2017). We estimated marginal trait means using r package emmeans (Lenth et al., 2025). Pairwise comparison of marginal means using Bonferroni correction for multiple comparisons was done as a post hoc test for difference in mean doubling times.

## Results

Replicate populations of *T. pyriformis* and *P. fluorescens* were evolved either alone or in co-culture at 26°C (close to the thermal optimum) or 30°C (a stressful temp) for 6 months (Figure 1A). Trait assays were then performed on all replicate populations (Figure 1B & C). We measured defensiveness in the bacterial prey and feeding efficiency in the ciliate predator, both traits that are expected evolve due to predator-prey interactions. We also measured growth rate, which is expected to evolve as a result of adaptation to temperature stress and to the laboratory environment. These traits were assayed for all populations at the final timepoint in the evolution experiment. Where possible, evolutionary trajectories were determined by assaying monthly timepoints from populations frozen during the course of the experiment.

### Temperature stress hinders the evolution of defense in the prey

Changes in defensiveness of bacteria were measured by assaying the ability of ancestral ciliates to divide when eating evolved vs. ancestral bacteria. Highly defended prey leads to low ciliate division rates. We found that prey defensiveness at the end of the evolution experiment depended on evolution condition. As expected, the presence of a ciliate predator resulted in increased defensiveness in bacterial populations (Figure 2A; GLMM, Type III Wald X^2^ (1) = 643.23, p= <2e-16). Defensiveness in bacteria evolved alone did not change whereas in populations evolved with predators defensiveness increased significantly (Figure 2A; One sample T-tests: 26°C alone t_15_ = -1.25, p=0.229; 30°C alone t_15_= 1.96, p=0.093; 26°C together t_15_ = 78.4, p= 2.05e-20; 30°C together t_15_= 35.4, p=1.44e-15). Although populations evolving with predators under both temperature regimes evolved increased defensiveness, populations evolving with the addition of temperature stress did not increase defensiveness as much as populations evolving at their optimum temperature, indicating that temperature stress hindered the evolution of defensiveness in these bacteria (Figure 2A; GLMM, Type III Wald X^2^ (1) = 78.88, p= <2e-16). Temperature stress, by itself, had no effect on the evolution of defensiveness (Figure 2A; GLMM, Type III Wald X^2^ (1) = 0.10, p= 0.75).

**Figure 2.**
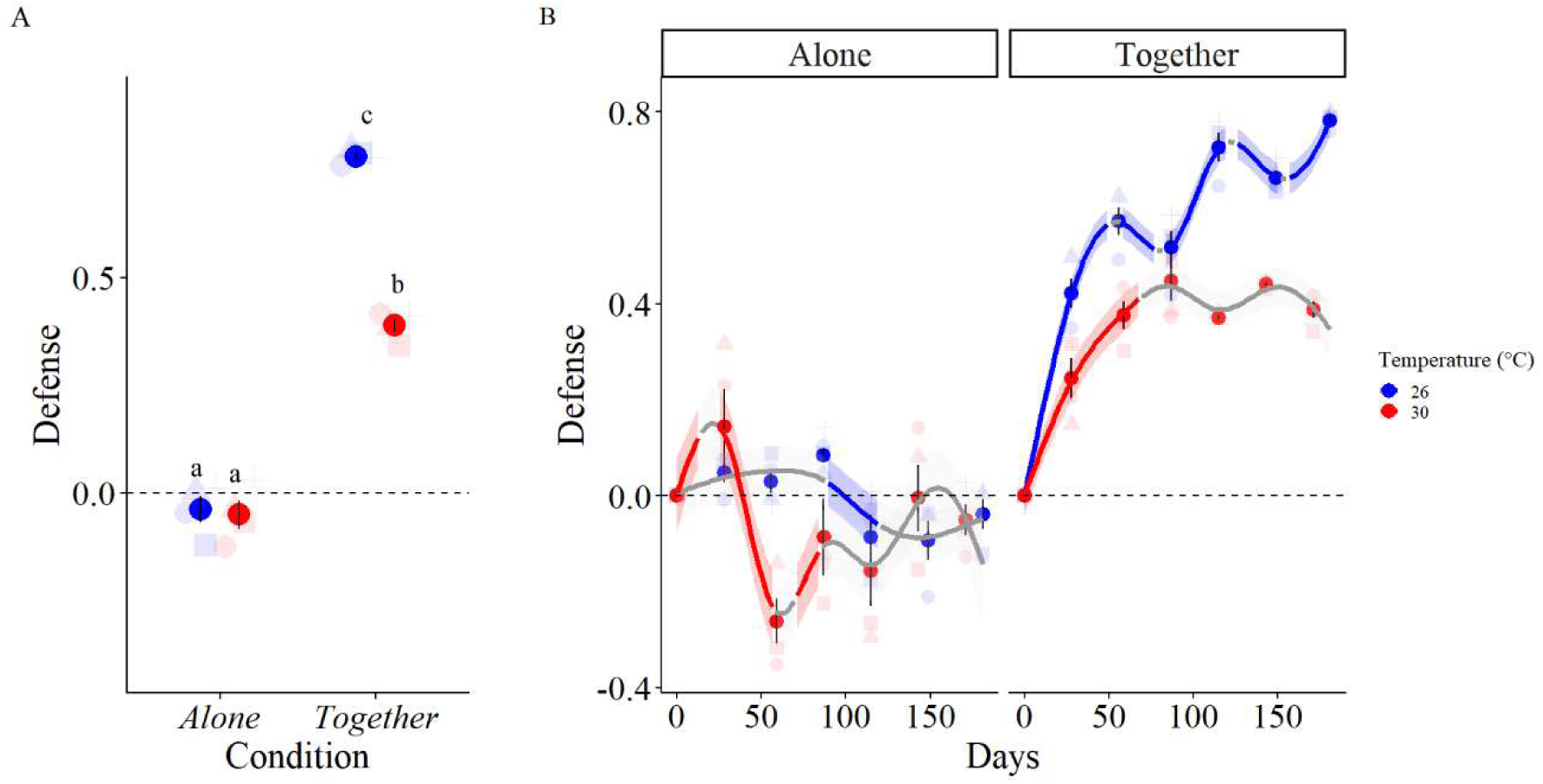
Evolution of defensiveness in bacterial prey is hindered by temperature stress. (A) Defensiveness of bacteria was measured at the final timepoint of the experimental evolution. Solid points correspond to the average trait value of the four replicate populations (faded dots) of an evolution condition. Color represents evolution temperature. Error bars indicate standard error of the mean. Letters denote groups that are significantly different based on Bonferroni-adjusted pairwise comparisons of estimated marginal means. Groups sharing letters do not differ. (B) Defense evolution trajectory over the entire course of the experimental evolution was measured by quantifying average defensiveness for populations from each month of the experiment. Panels represent trait trajectories for populations of bacteria evolving without (left) or with predators (right). Color represents evolution temperature. Ribbon represents the 95% confidence interval for the gam fit.

We then measured evolutionary trajectories of defensiveness in all populations by assaying defensiveness from samples that had been frozen each month of the evolution experiment. Trajectories were fit using GAM since trajectories were not linear (ΔAICc = 306.12, linear vs. smooth model). Trajectories for populations evolving in different conditions were different (ΔAICc = 385.25, universal smooth vs. smooth by group). Defense changed very little over the course of the experiment in populations evolving alone. However, alone populations at 30°C initially increased then decreased in defense, eventually returning to ancestral levels (Fig 2B). Defensiveness in populations evolving with predation all increased during the initial phase of adaptation, but populations evolving at 26°C increased in defensiveness faster than populations evolving under temperature stress (Fig 2B). After approximately day 60, defensiveness in 30°C populations stopped increasing, while defensiveness in populations evolving at the optimum temperature continued to increase overall though less rapidly and with occasional decreases (Fig 2B).

### Temperature stress hinders the evolution of predator feeding efficiency

Changes in ciliate predator feeding efficiency were quantified similarly to defense, in an assay that measured their ability to consume and use ancestral bacterial prey. At the end of the experimental evolution, we found that all populations, except for 26°C alone, evolved increased feeding efficiency compared to the ancestor (Figure 3; One sample T-tests: 26°C alone t_15_ = 1.44, p=0.172; 30°C alone t_15_= 2.38, p=0.041; 26°C together t_15_ = 10.3, p= 1.44e-8; 30°C together t_15_= 5.03, p=0.0003). Similar to defense evolution, the presence of bacterial prey significantly influenced feeding efficiency (Figure 2B; ANOVA: F_1,12_= 27.26, p= 0.0002). Temperature stress also affected the evolution of feeding efficiency when evolved with bacterial prey (Figure 3; ANOVA: F_1,12_= 7.45, p= 0.018). In particular, while feeding efficiency increased in the 30°C together populations, that increase was not significantly different from the alone populations, whereas the 26°C together populations showed a significant increase relative to all other populations (Figure 3; Tukey test). Temperature stress, by itself, had no effect on trait evolution (Figure 3; ANOVA: F_1,12_= 0.141, p=0.714). Evolutionary trajectories were unable to be measured for feeding efficiency.

**Figure 3.**
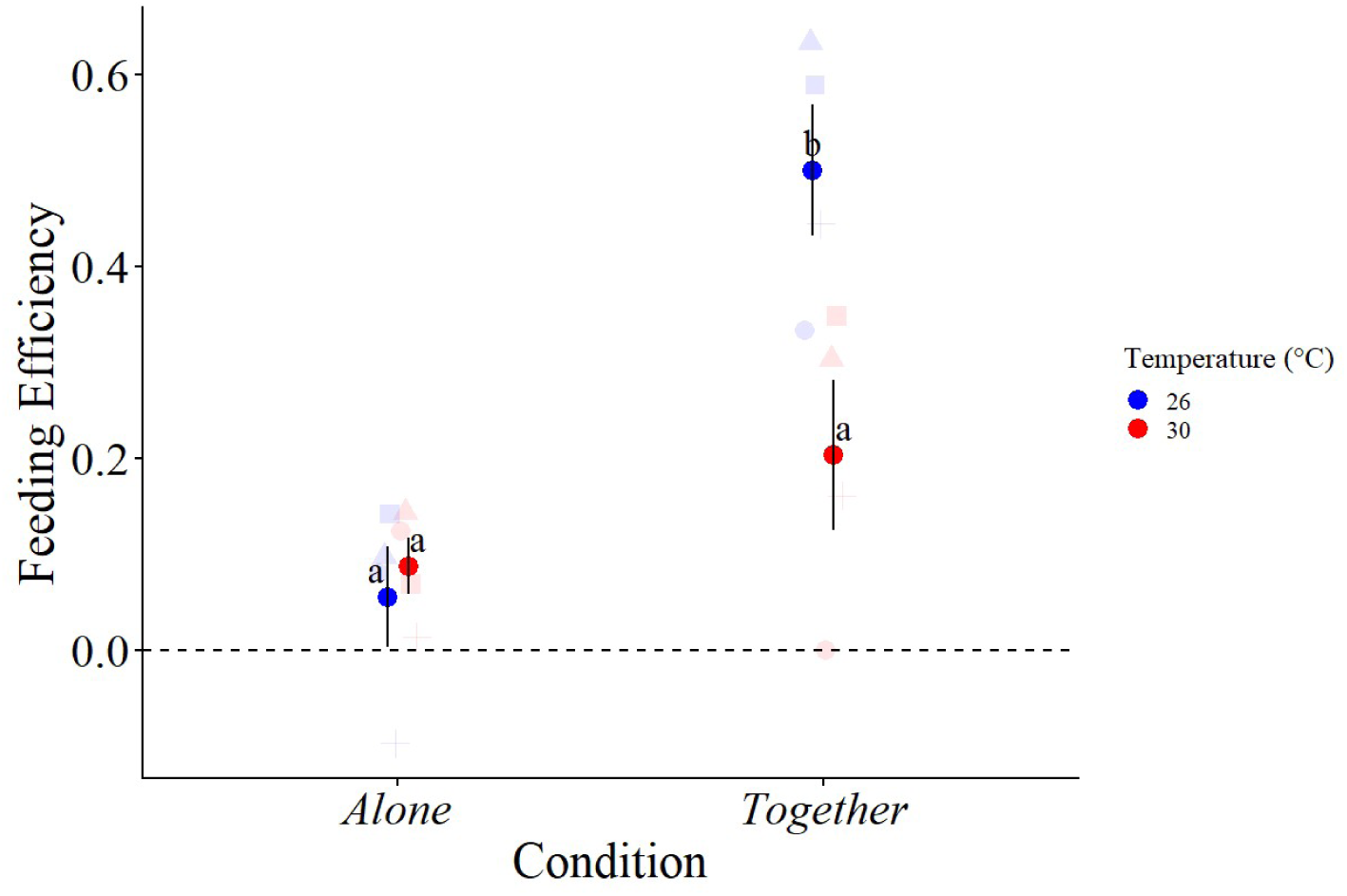
Evolution of feeding efficiency in ciliate predators is hindered by temperature stress. Feeding efficiency of ciliate predators was measured for populations at the final timepoint of the experimental evolution. Solid points correspond to the average trait value of the four replicate populations (faded dots). Trait values are shown for populations of ciliates evolving without (left) or with prey (right). Color represents evolution temperature. Error bars indicate standard error of the mean. Letters denote groups that are significantly different based on Bonferroni-adjusted pairwise comparisons of estimated marginal means. Groups sharing letters do not differ.

### Growth rates evolve in response to species interactions

Growth rates for both species were measured by generating growth curves axenically on growth medium, then estimating growth rate (divisions/hour) from curve fits. Growth rate evolution was quantified by measuring changes from ancestral growth rates over the course of the evolution experiment. There was no significant change in bacterial growth rates except for the 30°C population evolving with predation, which experienced transient changes in growth, with an overall decrease in growth rate over time (Figure 4). There was a small decrease in growth rate in populations evolved at 28°C, but the decreases are likely not biologically meaningful (Figure 4).

**Figure 4.**
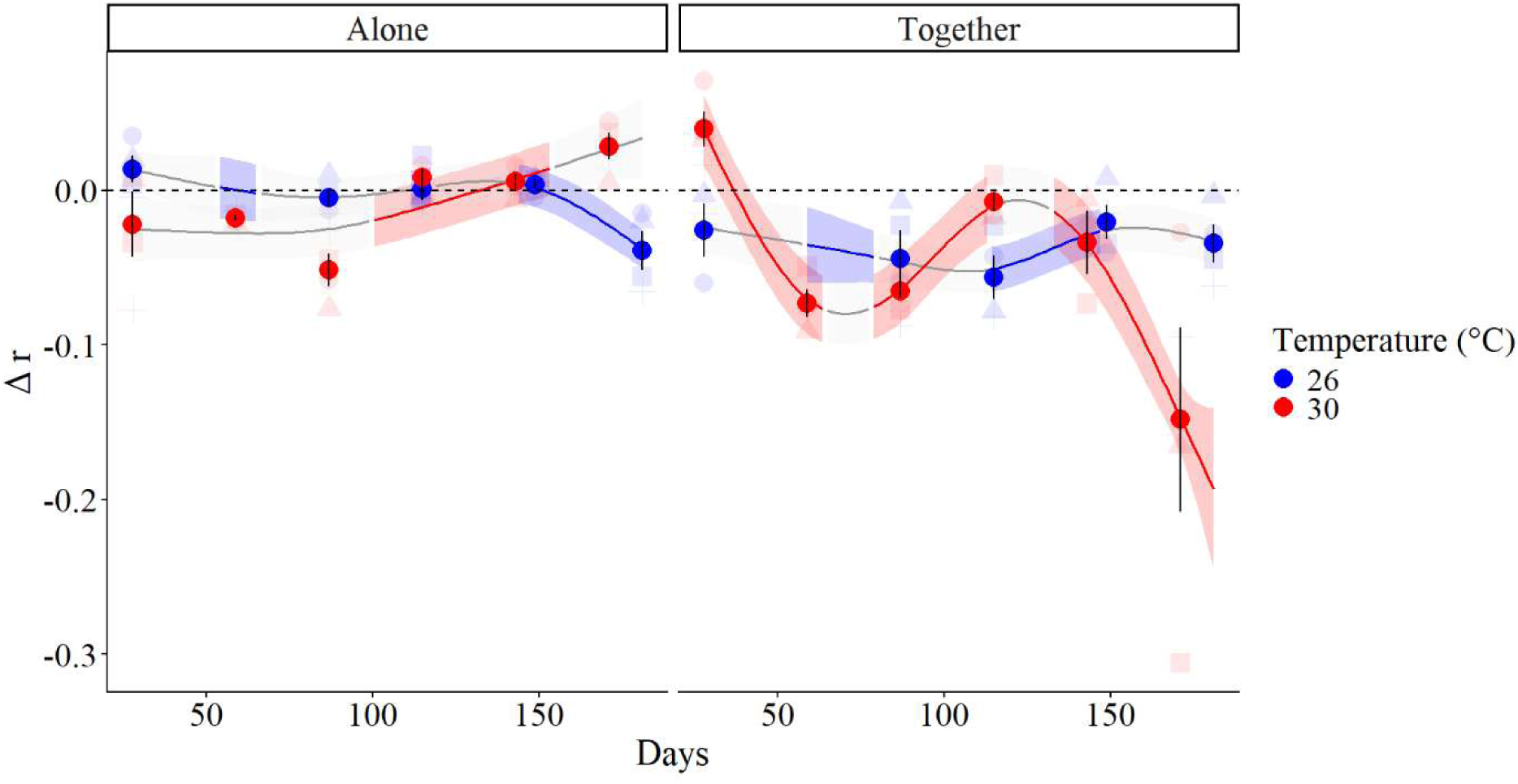
Growth rate of *P. fluorescens* decreases following evolution under temperature and predation stress. Solid points represent mean Δr (division/hour) of biological replicates of an evolution condition. Left panel shows the alone evolved populations; right panel shows the coevolved populations. Color represents evolution temperature. Ribbon represents the 95% confidence interval for the gam fit. Highlighted trends indicate periods of time where slope is different from 0, indicating periods of time where trait change was significant.

There was no change in growth rates for any group of ciliates when grown axenically on nutrient rich medium (Figure 5A). In the alone evolved populations, there were similar changes in growth rates between populations evolving at the two different temperatures, likely reflecting adaptation to laboratory conditions (Fig 5A). Populations evolving with prey at 28°C show similar trajectories to those evolving without prey, but with a more pronounced decrease during the first half of the experiment (Fig 5A). Populations evolving with prey at 30°C show a linear decline in growth rates (Fig 5A), but it is unclear whether a decrease of 0.025 divisions/per hour is likely to be biologically meaningful on these short time scales.

**Figure 5.**
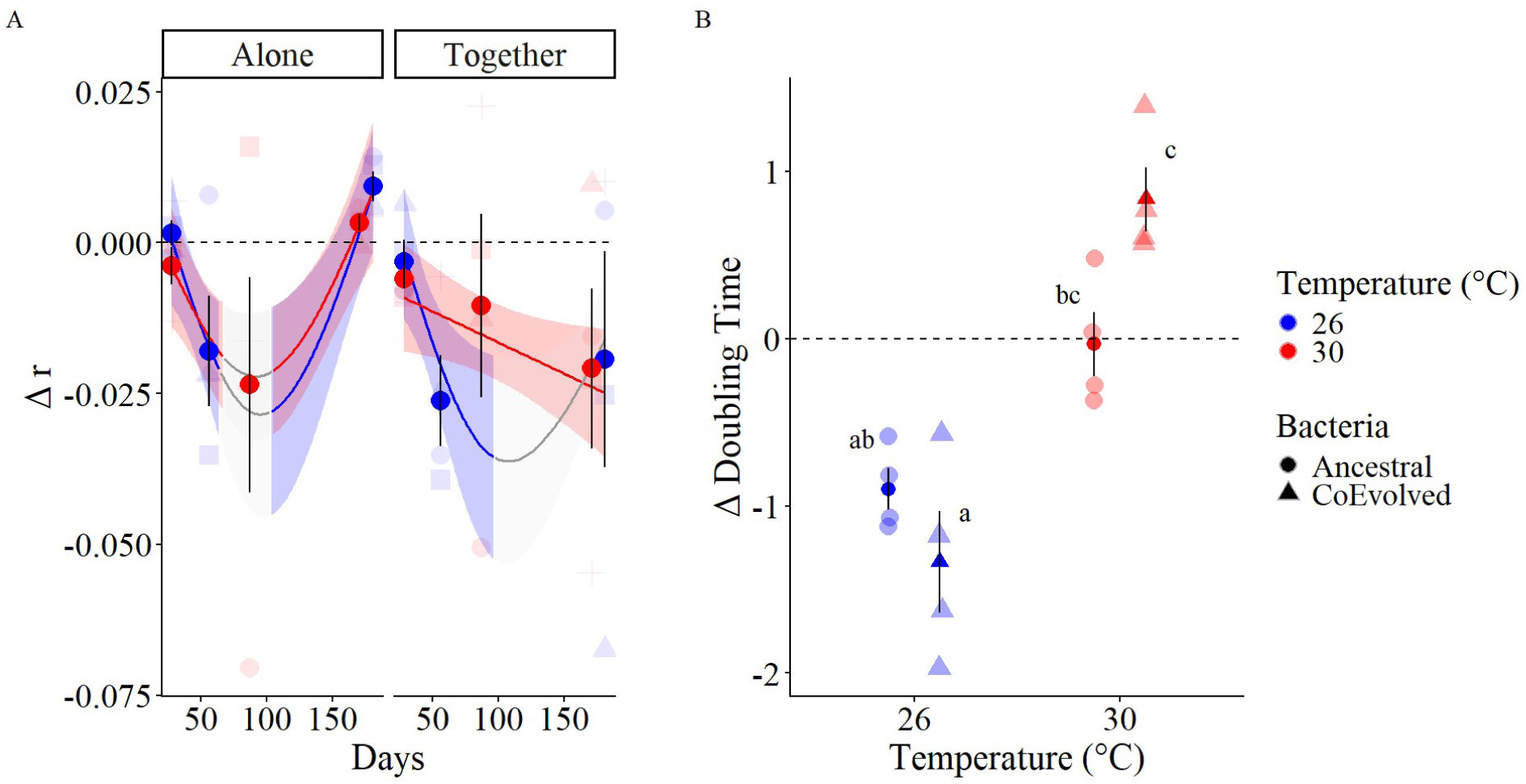
Growth rate of *T. pyriformis* on axenic medium shows little change, but growth on coevolving prey is hindered by temperature stress (A) Changes in ciliate growth rates in axenic medium. Solid points represent mean Δr (division/hour) of biological replicates of an evolution condition. Left panel shows the alone evolved populations; right panel shows the coevolved populations. Color represents evolution temperature. Ribbon represents the 95% confidence interval for the gam fit. Highlighted trends indicated periods of time where slope is different form 0, indicating periods of time where trait change was significant. (B) Ciliate growth depends on bacteria type. Mean doubling times (hours) of biological replicate populations of coevolved ciliates evolved at both temperatures on either co-evolved or ancestral bacteria. Color represents evolution temperature. Shape represents bacteria type. Error bars indicate standard error of the mean. Letters indicate significant differences between average trait value between evolution conditions.

Since populations of ciliate predators evolving together with bacterial prey were using bacteria as a food source as opposed to axenic medium, we also measured ciliate doubling times when consuming bacteria. We tested for changes in growth rates by measuring the doubling times of the coevolved ciliate populations on ancestral as well as coevolved bacteria. A decrease in doubling time indicates faster growth of the ciliate population. We found that temperature influenced doubling time evolution (Figure 5B; ANOVA Temperature F_1,12_= 7.90, p= 0.015), with 26°C populations showing a larger decrease in doubling time, but bacterial identity, by itself, had no effect on this trait (Figure 5B; ANOVA F_1,12_= 2.05, p= 0.177). Bacterial identity only affected doubling time evolution under temperature stress (Figure 5B; ANOVA F_1,12_= 9.13, p= 0.011). On average, populations evolving at 26°C evolved to grow faster than their ancestors when fed bacteria, and there is a trend toward growing faster on their coevolved bacteria than on ancestral bacteria (Figure 5B). Coevolved ciliate populations evolving under temperature stress were able to use ancestral bacteria to the same extent as their ciliate ancestors, but interestingly ancestral ciliates tended to do better on 30°C coevolved bacteria than their coevolved ciliate partners did (though the difference is not statically significant; Figure 5B).

## Discussion

In this study, we tested the effects of temperature stress on predator-prey coevolution using the microbial *T. pyriformis – P. fluorescens* system. Under our experimental conditions, we expected (i) defensiveness of bacterial prey to evolve in response to predation, (ii) feeding efficiency of ciliate predators to evolve in response to using bacteria as a food source, (iii) both of these traits to evolve more slowly under temperature stress, and (iv) growth rates of both species to change in response to selection under experimental conditions. As expected, we found that defensiveness and feeding efficiency did increase in coevolved populations and that the evolution of these traits was hindered under temperature stress. We did not observe the expected increases in growth rates usually seen as a response to increased temperature and adaptation to lab conditions. We did, however, observe a tradeoff between growth rates and defensiveness in coevolved bacteria populations, but interestingly that only manifested under temperature stress.

### Temperature stress slows coevolution

As predicted, coevolved bacterial populations evolved increased defensiveness over the course of the experiment. Defenses against predation in bacteria have been shown to include the formation of biofilms and the production of toxins (Hoque et al., 2023; Matz & Kjelleberg, 2005; Seiler et al., 2017). For example, under predation pressure, populations of *P. fluorescens* have been shown to evolve biofilm-forming morphotypes (Friman et al., 2014; Meyer & Kassen, 2007; Spiers et al., 2003). We do not know the mechanism of defense in our study, however, based on these previous studies and observation of our cultures, it is likely that biofilm formation is involved.

Importantly, consistent with our overarching hypothesis, we found that temperature stress hindered the evolution of defense in this system. At the optimum temperature, bacterial defense continued to increase throughout the experiment, consistent with an evolutionary arms race between predator and prey. In contrast, following a slower initial increase, defensiveness plateaued in the bacterial populations evolved at a stressful temperature, suggesting that temperature stress limits defense evolution in bacteria. Alternatively, this result could be due to decreased selection pressure from a predator unable to adapt under temperature stress. This result is similar to a result from Hiltunen et al. (2015) showing that fluctuations in salinity or nutrient stress hinder evolution of prey defense in a *T. thermophila – P. fluorescens* system; no effects were seen in the predator in that system, however.

Similar to defensiveness, as expected, the predator populations evolved increased feeding efficiency when coevolved with prey. Mechanisms of increased feeding efficiency in ciliates are less well studied, but changes in morphology and swimming patterns have been shown to be associated with changes in foraging behavior when feeding on evolving prey (Cairns et al., 2020). Also similar to defensiveness, populations evolved under temperature stress evolved significantly lower levels of feeding efficiency than those evolved at the thermal optimum. This effect in the predators seems to be even stronger than in the prey, with the 26°C predator populations having over double the feeding efficiency of the 30°C populations.

There are several possible reasons that temperature stress might hinder the evolution of traits involved in species interactions. First, growth rates slowed under temperature stress in both species. This would result in fewer generations of selection in the stressed populations. For example, we estimate that in 6 months, ∼768 generations passed in the 26°C bacteria populations, but only ∼508 generation in populations evolving with predation at 30°C. Population sizes could also be reduced under stress, resulting in a decreased supply of mutations and increased genetic drift. Temperature stress has also been shown to reshape predator-prey ecology by altering attack rates, handling times, metabolism, and mortality. Models have shown that warming can also change the amplitude and period of population cycles, and, near thermal limits, erode top-down control when predators are disproportionately affected (DeLong & Lyon, 2020).

### Growth rates tradeoff with defense and feeding efficiency

Microbial growth rates are influenced by environmental factors, such as resource type, temperature, and biotic interactions (Gonzalez & Aranda, 2023; Iven et al., 2022). In any given environment, growth rates tend to increase with increasing temperature until an optimum is reached, above which point growth rates rapidly decline, following a stereotypical unimodal thermal performance curve (Huey & Kingsolver, 1993; Malusare et al., 2023).

In microbial experimental evolution, growth rates normally increase over time due to adaptation to laboratory conditions (Dragosits & Mattanovich, 2013; Hirasawa & Maeda, 2023; Knöppel et al., 2018; McDonald, 2019; Tarkington & Zufall, 2021; Wiser et al., 2013). Microbial populations readily adapt to environmental stressors. Both bacterial and ciliate systems have demonstrated that populations can evolve increased growth ability following an initial fitness decline due to temperature stress (Bennett et al., 1992; Bennett & Lenski, 1993; Duncan et al., 2011). Surprisingly, we did not observe increased growth rates in either species evolved alone (Fig. 4 and 5A). One possible explanation may come from our transfer regime, which resulted in populations experiencing long periods of time in stationary phase. This could lead to strong selection for increases in intraspecific competition for resources rather than growth rate.

The pattern of growth rate evolution in the coevolving populations was more complicated. At 26°C, neither species showed increased growth rates when cultured axenically. At 30°C, coevolved populations of both species showed significant declines in growth rate (whether grown axenically or fed prey). The only condition under which we observed increased growth rates was in coevolved ciliate populations at 26°C and fed ancestral bacteria. This suggests that in the absence of temperature stress, ciliates are able to evolve general adaptations to consuming bacteria (since these populations showed increased growth rates when fed either ancestral or evolved bacteria).

The overall lack of evolved increases in growth rate in coevolving populations may be explained with reference to previous studies that have demonstrated that under antagonistic coevolution, evolution of defense traits can tradeoff with growth rate (Ehrlich et al., 2020; Friman et al., 2014; Guillonneau et al., 2022; Lürling, 2021; Pančić & Kiørboe, 2018; Yoshida et al., 2004). For example, a study of a rotifer-alga predator-prey system found that algae that were more defensive against rotifer predators had lower growth rates and diminished competitive ability. Additionally, these tradeoffs can be amplified with the addition of environmental stress (O’Donnell et al., 2013; Zhu et al., 2016). It seems likely, then, that tradeoffs in the response to selection on defense and feeding traits, amplified at the higher temperature, may be responsible for our unexpected growth rate results. Further studies are needed to address this possibility.

### Summary

By studying how an environmental stress jointly affects both predator and prey evolution, this study reveals that temperature stress hinders the evolution of traits important in species interactions and may amplify the tradeoffs between these traits and fitness-associated traits such as growth rate. These results suggest that population dynamics of coevolving species may be less stable due to limited ability to co-adapt under increasing warming—a potential threat to ecosystem stability under climate change. Future studies should explicitly measure eco-evolutionary dynamics and consider conditions more similar to those experienced in nature, such as additional interacting species and fluctuating temperature stress.

## References

Abrams, P. A. (2000). The evolution of predator-prey interactions: Theory and evidence. Annual Review of Ecology, Evolution, and Systematics, 31(Volume 31, 2000), 79–105. 10.1146/annurev.ecolsys.31.1.79

Abrams, P. A., & Matsuda, H. (1996). Fitness minimization and dynamic instability as a consequence of predator-prey coevolution. Evolutionary Ecology, 10(2), 167–186. 10.1007/BF01241783

Bennett, A. F., & Hughes, B. S. (2009). Microbial experimental evolution. *American Journal of Physiology-Regulatory*, Integrative and Comparative Physiology, 297(1), R17–R25. 10.1152/ajpregu.90562.2008

Bennett, A. F., & Lenski, R. E. (1993). Evolutionary Adaptation to Temperature Ii. Thermal Niches of Experimental Lines of Escherichia Coli. Evolution, 47(1), 1–12. 10.1111/j.1558-5646.1993.tb01194.x

Bennett, A. F., Lenski, R. E., & Mittler, J. E. (1992). Evolutionary adaptation to temperature. I. Fitness responses of escherichia coli to changes in its thermal environment. Evolution, 46(1), 16–30. 10.1111/j.1558-5646.1992.tb01981.x

Borghans, J. A. M., Beltman, J. B., & De Boer, R. J. (2004). MHC polymorphism under host-pathogen coevolution. Immunogenetics, 55(11), 732–739. 10.1007/s00251-003-0630-5

Brockhurst, M. A., & Koskella, B. (2013). Experimental coevolution of species interactions. Trends in Ecology & Evolution, 28(6), 367–375. 10.1016/j.tree.2013.02.009

Brooks, M., Kristensen, K., Benthem, K. van, Magnusson, A., Berg, C., Nielsen, A., Skaug, H., Mächler, M., & Bolker, B. (2017). glmmTMB Balances Speed and Flexibility Among Packages for Zero-inflated Generalized Linear Mixed Modeling. The R Journal. https://digitalcommons.unl.edu/r-journal/675

Buckling, A., & Rainey, P. B. (2002). Antagonistic coevolution between a bacterium and a bacteriophage. *Proceedings*. Biological Sciences, 269(1494), 931–936. 10.1098/rspb.2001.1945

Burak, M. K., Monk, J. D., & Schmitz, O. J. (2018). Eco-evolutionary dynamics: The predator-prey adaptive play and the ecological theater. The Yale Journal of Biology and Medicine, 91(4), 481–489.

Cairns, J., Moerman, F., Fronhofer, E. A., Altermatt, F., & Hiltunen, T. (2020). Evolution in interacting species alters predator life-history traits, behaviour and morphology in experimental microbial communities. Proceedings of the Royal Society B: Biological Sciences, 287(1928), 20200652. 10.1098/rspb.2020.0652

Cassidy-Hanley, D. M. (2012). Chapter 8 - *Tetrahymena* in the laboratory: Strain resources, methods for culture, maintenance, and storage. In K. Collins (Ed.), Methods in Cell Biology (Vol. 109, pp. 237–276). Academic Press. 10.1016/B978-0-12-385967-9.00008-6

Cavicchioli, R., Ripple, W. J., Timmis, K. N., Azam, F., Bakken, L. R., Baylis, M., Behrenfeld, M. J., Boetius, A., Boyd, P. W., Classen, A. T., Crowther, T. W., Danovaro, R., Foreman, C. M., Huisman, J., Hutchins, D. A., Jansson, J. K., Karl, D. M., Koskella, B., Mark Welch, D. B., … Webster, N. S. (2019). Scientists’ warning to humanity: Microorganisms and climate change. Nature Reviews Microbiology, 17(9), 569–586. 10.1038/s41579-019-0222-5

Dawkins, R., & Krebs, J. R. (1997). Arms races between and within species. Proceedings of the Royal Society of London. Series B. Biological Sciences, 205(1161), 489–511. 10.1098/rspb.1979.0081

DeLong, J. P., & Lyon, S. (2020). Temperature alters the shape of predator-prey cycles through effects on underlying mechanisms. PeerJ, 8, e9377. 10.7717/peerj.9377

Dieckmann, U., Marrow, P., & Law, R. (1995). Evolutionary cycling in predator-prey interactions: Population dynamics and the red queen. Journal of Theoretical Biology, 176(1), 91–102. 10.1006/jtbi.1995.0179

Doerder, F. P. (2019). Barcodes Reveal 48 New species of *Tetrahymena*, *Dexiostoma*, and *Glaucoma*: Phylogeny, ecology, and biogeography of new and established species. Journal of Eukaryotic Microbiology, 66(1), 182–208. 10.1111/jeu.12642

Doerder, P., & Brunk, C. (2012). Natural populations and inbred strains of *Tetrahymena*. In Methods in Cell Biology (Vol. 109, pp. 277–300). Academic Press. 10.1016/B978-0-12-385967-9.00009-8

Dragosits, M., & Mattanovich, D. (2013). Adaptive laboratory evolution – principles and applications for biotechnology. Microbial Cell Factories, 12, 64. 10.1186/1475-2859-12-64

Draper, A. M., & Weissburg, M. J. (2019). Impacts of global warming and elevated co2 on sensory behavior in predator-prey interactions: a review and synthesis. Frontiers in Ecology and Evolution, 7. 10.3389/fevo.2019.00072

Duncan, A. B., Fellous, S., Quillery, E., & Kaltz, O. (2011). Adaptation of *Paramecium caudatum* to variable conditions of temperature stress. Research in Microbiology, 162(9), 939–944. 10.1016/j.resmic.2011.04.012

Ehrlich, E., Kath, N. J., & Gaedke, U. (2020). The shape of a defense-growth trade-off governs seasonal trait dynamics in natural phytoplankton. The ISME Journal, 14(6), 1451–1462. 10.1038/s41396-020-0619-1

Eisenmann, H., Harms, H., Meckenstock, R., Meyer, E. I., & Zehnder, A. J. B. (1998). Grazing of a *Tetrahymena* sp. on adhered bacteria in percolated columns monitored by in situ hybridization with fluorescent oligonucleotide probes. Applied and Environmental Microbiology, 64(4), 1264–1269. 10.1128/aem.64.4.1264-1269.1998

Finlay, B. J., & Esteban, G. F. (1998). Freshwater protozoa: Biodiversity and ecological function. Biodiversity and Conservation, 7(9), 1163–1186. 10.1023/A:1008879616066

Fox, J., Weisberg, S., Price, B., Adler, D., Bates, D., Baud-Bovy, G., Bolker, B., Ellison, S., Firth, D., Friendly, M., Gorjanc, G., Graves, S., Heiberger, R., Krivitsky, P., Laboissiere, R., Maechler, M., Monette, G., Murdoch, D., Nilsson, H., Ogle, D., Ripley, B., Short, T., Venables, W., Walker, S., Winsemius, D., Zeileis, A., R-Core. (2024). car: Companion to Applied Regression (Version 3.1-3) [Computer software]. https://cran.r-project.org/web/packages/car/index.html

Friman, V.-P., Hiltunen, T., Laakso, J., & Kaitala, V. (2008). Availability of prey resources drives evolution of predator–prey interaction. Proceedings of the Royal Society B: Biological Sciences, 275(1643), 1625–1633. 10.1098/rspb.2008.0174

Friman, V.-P., Jousset, A., & Buckling, A. (2014). Rapid prey evolution can alter the structure of predator–prey communities. Journal of Evolutionary Biology, 27(2), 374–380. 10.1111/jeb.12303

Fussmann, G. F., Ellner, S. P., Shertzer, K. W., & Hairston Jr., N. G. (2000). Crossing the Hopf bifurcation in a live predator-prey system. Science, 290(5495), 1358–1360. 10.1126/science.290.5495.1358

Getz, W. M., & Owen-Smith, N. (2011). Consumer-resource dynamics: quantity, quality, and allocation. PLOS ONE, 6(1), e14539. 10.1371/journal.pone.0014539

Gonzalez, J. M., & Aranda, B. (2023). Microbial Growth under Limiting Conditions-Future Perspectives. Microorganisms, 11(7), 1641. 10.3390/microorganisms11071641

Gorovsky, M. A., Yao, M.-C., Keevert, J. B., & Pleger, G. L. (1975). Chapter 16 isolation of micro- and macronuclei of *Tetrahymena pyriformis*. In D. M. Prescott (Ed.), Methods in Cell Biology (Vol. 9, pp. 311–327). Academic Press. 10.1016/S0091-679X(08)60080-1

Gougoulias, C., Clark, J. M., & Shaw, L. J. (2014). The role of soil microbes in the global carbon cycle: Tracking the below-ground microbial processing of plant-derived carbon for manipulating carbon dynamics in agricultural systems. Journal of the Science of Food and Agriculture, 94(12), 2362–2371. 10.1002/jsfa.6577

Govaert, L., Fronhofer, E. A., Lion, S., Eizaguirre, C., Bonte, D., Egas, M., Hendry, A. P., De Brito Martins, A., Melián, C. J., Raeymaekers, J. A. M., Ratikainen, I. I., Saether, B.-E., Schweitzer, J. A., & Matthews, B. (2019). Eco-evolutionary feedbacks—Theoretical models and perspectives. Functional Ecology, 33(1), 13–30. 10.1111/1365-2435.13241

Guillonneau, R., Murphy, A. R. J., Teng, Z.-J., Wang, P., Zhang, Y.-Z., Scanlan, D. J., & Chen, Y. (2022). Trade-offs of lipid remodeling in a marine predator–prey interaction in response to phosphorus limitation. Proceedings of the National Academy of Sciences, 119(36), e2203057119. 10.1073/pnas.2203057119

Gupta, A., Gupta, R., & Singh, R. L. (2016). Microbes and Environment. Principles and Applications of Environmental Biotechnology for a Sustainable Future, 43–84. 10.1007/978-981-10-1866-4_3

Harrison, E., Laine, A.-L., Hietala, M., & Brockhurst, M. A. (2013). Rapidly fluctuating environments constrain coevolutionary arms races by impeding selective sweeps. Proceedings of the Royal Society B: Biological Sciences, 280(1764), 20130937. 10.1098/rspb.2013.0937

Hartig, F., Lohse, L., & leite, M. de S. (2024). DHARMa: Residual Diagnostics for Hierarchical (Multi-Level / Mixed) Regression Models (Version 0.4.7) [Computer software]. https://cran.r-project.org/web/packages/DHARMa/index.html

Hiltunen, T., Ayan, G. B., & Becks, L. (2015). Environmental fluctuations restrict eco-evolutionary dynamics in predator–prey system. Proceedings of the Royal Society B: Biological Sciences, 282(1808), 20150013. 10.1098/rspb.2015.0013

Hiltunen, T., & Becks, L. (2014). Consumer co-evolution as an important component of the eco-evolutionary feedback. Nature Communications, 5(1), 5226. 10.1038/ncomms6226

Hiltunen, T., Cairns, J., Frickel, J., Jalasvuori, M., Laakso, J., Kaitala, V., Künzel, S., Karakoc, E., & Becks, L. (2018). Dual-stressor selection alters eco-evolutionary dynamics in experimental communities. Nature Ecology & Evolution, 2(12), Article 12. 10.1038/s41559-018-0701-5

Hirasawa, T., & Maeda, T. (2023). Adaptive Laboratory Evolution of Microorganisms: Methodology and Application for Bioproduction. Microorganisms, 11(1), 92. 10.3390/microorganisms11010092

Hochberg, M. E., & Baalen, M. van. (1998). Antagonistic coevolution over productivity Gradients. The American Naturalist, 152(4), 620–634. 10.1086/286194

Hogle, S. L., Ruusulehto, L., Cairns, J., Hultman, J., & Hiltunen, T. (2023). Localized coevolution between microbial predator and prey alters community-wide gene expression and ecosystem function. The ISME Journal, 17(4), 514–524. 10.1038/s41396-023-01361-9

Hoque, M. M., Espinoza-Vergara, G., & McDougald, D. (2023). Protozoan predation as a driver of diversity and virulence in bacterial biofilms. FEMS Microbiology Reviews, 47(4), fuad040. 10.1093/femsre/fuad040

Huey, R. B., & Kingsolver, J. G. (1993). Evolution of resistance to high temperature in ectotherms. The American Naturalist, 142, S21–S46. 10.1086/285521

Iven, H., Walker, T. W. N., & Anthony, M. (2022). Biotic Interactions in Soil are Underestimated Drivers of Microbial Carbon Use Efficiency. Current Microbiology, 80(1), 13. 10.1007/s00284-022-02979-2

Jack, J. D., & Gilbert, J. J. (1997). Effects of Metazoan Predators on Ciliates in Freshwater Plankton Communities1. Journal of Eukaryotic Microbiology, 44(3), 194–199. 10.1111/j.1550-7408.1997.tb05699.x

Janssens, L., Van Dievel, M., & Stoks, R. (2015). Warming reinforces nonconsumptive predator effects on prey growth, physiology, and body stoichiometry. Ecology, 96(12), 3270–3280. 10.1890/15-0030.1

Jassey, V. E. J., Walcker, R., Kardol, P., Geisen, S., Heger, T., Lamentowicz, M., Hamard, S., & Lara, E. (2022). Contribution of soil algae to the global carbon cycle. New Phytologist, 234(1), 64–76. 10.1111/nph.17950

Jousset, A., Lara, E., Wall, L. G., & Valverde, C. (2006). Secondary metabolites help biocontrol strain *Pseudomonas fluorescens* CHA0 to escape protozoan grazing. Applied and Environmental Microbiology, 72(11), 7083–7090. 10.1128/AEM.00557-06

Jousset, A., Rochat, L., Péchy-Tarr, M., Keel, C., Scheu, S., & Bonkowski, M. (2009). Predators promote defence of rhizosphere bacterial populations by selective feeding on non-toxic cheaters. The ISME Journal, 3(6), 666–674. 10.1038/ismej.2009.26

Kawecki, T. J., Lenski, R. E., Ebert, D., Hollis, B., Olivieri, I., & Whitlock, M. C. (2012). Experimental evolution. Trends in Ecology & Evolution, 27(10), 547–560. 10.1016/j.tree.2012.06.001

Kingsland, S. (2015). Alfred J. Lotka and the origins of theoretical population ecology. Proceedings of the National Academy of Sciences of the United States of America, 112(31), 9493–9495. 10.1073/pnas.1512317112

Knöppel, A., Knopp, M., Albrecht, L. M., Lundin, E., Lustig, U., Näsvall, J., & Andersson, D. I. (2018). Genetic Adaptation to Growth Under Laboratory Conditions in *Escherichia coli* and *Salmonella enterica*. Frontiers in Microbiology, 9. 10.3389/fmicb.2018.00756

Kuznetsova, A., Brockhoff, P. B., & Christensen, R. H. B. (2017). lmerTest Package: Tests in Linear Mixed Effects Models. Journal of Statistical Software, 82, 1–26. 10.18637/jss.v082.i13

Lenth, R. V., Piaskowski, J., Banfai, B., Bolker, B., Buerkner, P., Giné-Vázquez, I., Hervé, M., Jung, M., Love, J., Miguez, F., Riebl, H., & Singmann, H. (2025). emmeans: Estimated Marginal Means, aka Least-Squares Means (Version 2.0.0) [Computer software]. https://cran.r-project.org/web/packages/emmeans/index.html?utm_source=chatgpt.com

Lürling, M. (2021). Grazing resistance in phytoplankton. Hydrobiologia, 848(1), 237–249. 10.1007/s10750-020-04370-3

Malusare, S. P., Zilio, G., & Fronhofer, E. A. (2023). Evolution of thermal performance curves: A meta-analysis of selection experiments. Journal of Evolutionary Biology, 36(1), 15–28. 10.1111/jeb.14087

Matz, C., & Kjelleberg, S. (2005). Off the hook – how bacteria survive protozoan grazing. Trends in Microbiology, 13(7), 302–307. 10.1016/j.tim.2005.05.009

McDonald, M. J. (2019). Microbial experimental evolution – a proving ground for evolutionary theory and a tool for discovery. EMBO Reports, 20(8), e46992. 10.15252/embr.201846992

Mei, S., Wang, M., Salles, J. F., & Hackl, T. (2024). Diverse rhizosphere-associated Pseudomonas genomes from along a Wadden Island salt marsh transition zone. Scientific Data, 11, 1140. 10.1038/s41597-024-03961-2

Meyer, J. R., & Kassen, R. (2007). The effects of competition and predation on diversification in a model adaptive radiation. Nature, 446(7134), 432–435. 10.1038/nature05599

Monod, J. (1949). The growth of bacterial cultures. Annual Review of Microbiology, 3(Volume 3, 1949), 371–394. 10.1146/annurev.mi.03.100149.002103

Nelder, J. A. (1961). The fitting of a generalization of the logistic curve. Biometrics, 17(1), 89–110. 10.2307/2527498

Northfield, T. D., & Ives, A. R. (2013). Coevolution and the effects of climate change on interacting species. PLOS Biology, 11(10), e1001685. 10.1371/journal.pbio.1001685

Nuismer, S. L., & Thompson, J. N. (2006). Coevolutionary alternation in antagonistic interactions. Evolution, 60(11), 2207–2217. 10.1111/j.0014-3820.2006.tb01858.x

O’Donnell, D. R., Fey, S. B., & Cottingham, K. L. (2013). Nutrient availability influences kairomone-induced defenses in *Scenedesmus acutus* (Chlorophyceae). Journal of Plankton Research, 35(1), 191–200. 10.1093/plankt/fbs083

Öhlund, G., Hedström, P., Norman, S., Hein, C. L., & Englund, G. (2015). Temperature dependence of predation depends on the relative performance of predators and prey. Proceedings of the Royal Society B: Biological Sciences, 282(1799), 20142254. 10.1098/rspb.2014.2254

Paine, R. T. (1966). Food web complexity and species diversity. The American Naturalist, 100(910), 65–75.

Pančić, M., & Kiørboe, T. (2018). Phytoplankton defence mechanisms: Traits and trade-offs. Biological Reviews, 93(2), 1269–1303. 10.1111/brv.12395

Pandey, M. (2023). Impact of aquatic microbes in nutrient cycling and energy flow in ecosystems. Journal of Marine Biology & Oceanography, 2023. https://www.scitechnol.com/peer-review/impact-of-aquatic-microbes-in-nutrient-cycling-and-energy-flow-in-ecosystems-vHXi.php?article_id=23232

Petzoldt, T. (2025). growthrates: Estimate Growth Rates from Experimental Data (Version 0.8.5) [Computer software]. https://cran.r-project.org/web/packages/growthrates/index.html

Plum, K., Tarkington, J., & Zufall, R. A. (2022). Experimental evolution in tetrahymena. Microorganisms, 10(2), 414. 10.3390/microorganisms10020414

Porter, K. G., Pace, M. L., & Battey, J. F. (1979). Ciliate protozoans as links in freshwater planktonic food chains. Nature, 277(5697), 563–565. 10.1038/277563a0

Rainey, P. B., & Travisano, M. (1998). Adaptive radiation in a heterogeneous environment. Nature, 394(6688), Article 6688. 10.1038/27900

Redford, K. H. (2023). Extending conservation to include Earth’s microbiome. Conservation Biology, 37(3), e14088. 10.1111/cobi.14088

Rhodes, M. E. (1959). The characterization of *Pseudomonas fluorescens*. Journal of General Microbiology, 21(1), 221–263. 10.1099/00221287-21-1-221

Robertson, M. L., & Hammill, E. (2021). Temperature and prey morphology influence attack rate and handling time in a predator–prey interaction. Hydrobiologia, 848(19), 4637–4646. 10.1007/s10750-021-04666-y

Scheuerl, T., Cairns, J., Becks, L., & Hiltunen, T. (2019). Predator coevolution and prey trait variability determine species coexistence. Proceedings of the Royal Society B: Biological Sciences, 286(1902), 20190245. 10.1098/rspb.2019.0245

Schmitz, O. (2017). Predator and prey functional traits: Understanding the adaptive machinery driving predator–prey interactions. F1000Research, 6, 1767. 10.12688/f1000research.11813.1

Seger, J. (1988). Dynamics of some simple host-parasite models with more than two genotypes in each species. *Philosophical Transactions of the Royal Society of London. B*, Biological Sciences, 319(1196), 541–555. 10.1098/rstb.1988.0064

Seiler, C., van Velzen, E., Neu, T. R., Gaedke, U., Berendonk, T. U., & Weitere, M. (2017). Grazing resistance of bacterial biofilms: A matter of predators’ feeding trait. FEMS Microbiology Ecology, 93(9), fix112. 10.1093/femsec/fix112

Shalev, O., Ashkenazy, H., Neumann, M., & Weigel, D. (2022). Commensal *Pseudomonas* protect *Arabidopsis thaliana* from a coexisting pathogen via multiple lineage-dependent mechanisms. The ISME Journal, 16(5), 1235–1244. 10.1038/s41396-021-01168-6

Sherr, E. B., & Sherr, B. F. (2002). Significance of predation by protists in aquatic microbial food webs. Antonie van Leeuwenhoek, 81(1), 293–308. 10.1023/A:1020591307260

Silby, M. W., Winstanley, C., Godfrey, S. A. C., Levy, S. B., & Jackson, R. W. (2011). *Pseudomonas* genomes: Diverse and adaptable. FEMS Microbiology Reviews, 35(4), 652–680. 10.1111/j.1574-6976.2011.00269.x

Simon, E. M., Nanney, D. L., & Doerder, F. P. (2008). The “*Tetrahymena pyriformis*” complex of cryptic species. Biodiversity and Conservation, 17(2), 365–380. 10.1007/s10531-007-9255-6

Simpson, G. L. (2018). Modelling palaeoecological time series using generalised additive models. Frontiers in Ecology and Evolution, 6. 10.3389/fevo.2018.00149

Simpson, G. L. (2024). gratia: An R package for exploring generalized additive models. Journal of Open Source Software, 9(104), 6962. 10.21105/joss.06962

Spiers, A. J., Bohannon, J., Gehrig, S. M., & Rainey, P. B. (2003). Biofilm formation at the air–liquid interface by the *Pseudomonas fluorescens* SBW25 wrinkly spreader requires an acetylated form of cellulose. Molecular Microbiology, 50(1), 15–27. 10.1046/j.1365-2958.2003.03670.x

Tarkington, J., & Zufall, R. A. (2021). Temperature affects the repeatability of evolution in the microbial eukaryote *Tetrahymena thermophila*. Ecology and Evolution, 11(19), 13139–13152. 10.1002/ece3.8036

Theodosiou, L., Hiltunen, T., & Becks, L. (2019). The role of stressors in altering eco-evolutionary dynamics. Functional Ecology, 33(1), 73–83. 10.1111/1365-2435.13263

Thompson, J. N. (2005). Coevolution: The geographic mosaic of coevolutionary arms races. Current Biology: CB, 15(24), R992–994. 10.1016/j.cub.2005.11.046

Thurman, J., Drinkall, J., & Parry, J. D. (2010). Digestion of bacteria by the freshwater ciliate *Tetrahymena pyriformis*. Aquatic Microbial Ecology, 60, 163–174. 10.3354/ame01413

Tseng, M., & O’Connor, M. I. (2015). Predators modify the evolutionary response of prey to temperature change. Biology Letters, 11(12), 20150798. 10.1098/rsbl.2015.0798

Tylianakis, J. M., Didham, R. K., Bascompte, J., & Wardle, D. A. (2008). Global change and species interactions in terrestrial ecosystems. Ecology Letters, 11(12), 1351–1363. 10.1111/j.1461-0248.2008.01250.x

Vander Zanden, M. J., & Vadeboncoeur, Y. (2002). Fishes as integrators of benthic and pelagic food webs in lakes. Ecology, 83(8), 2152–2161. 10.1890/0012-9658(2002)083%255B2152:FAIOBA%255D2.0.CO;2

Walker, R., Wilder, S. M., & González, A. L. (2020). Temperature dependency of predation: Increased killing rates and prey mass consumption by predators with warming. Ecology and Evolution, 10(18), 9696–9706. 10.1002/ece3.6581

Weitere, M., Bergfeld, T., Rice, S. A., Matz, C., & Kjelleberg, S. (2005). Grazing resistance of *Pseudomonas aeruginosa* biofilms depends on type of protective mechanism, developmental stage and protozoan feeding mode. Environmental Microbiology, 7(10), 1593–1601. 10.1111/j.1462-2920.2005.00851.x

Wiser, M. J., Ribeck, N., & Lenski, R. E. (2013). Long-term dynamics of adaptation in asexual populations. Science, 342(6164), 1364–1367. 10.1126/science.1243357

Wood, S. N. (2017). Generalized Additive Models: An Introduction with R, Second Edition (2nd ed.). Chapman and Hall/CRC. 10.1201/9781315370279

Yoshida, T., Hairston, N. G., & Ellner, S. P. (2004). Evolutionary trade–off between defence against grazing and competitive ability in a simple unicellular alga, *Chlorella vulgaris*. Proceedings of the Royal Society of London. Series B: Biological Sciences, 271(1551), 1947–1953. 10.1098/rspb.2004.2818

Zhou, W., Qi, D., Swaisgood, R. R., Wang, L., Jin, Y., Wu, Q., Wei, F., & Nie, Y. (2021). Symbiotic bacteria mediate volatile chemical signal synthesis in a large solitary mammal species. The ISME Journal, 15(7), 2070–2080. 10.1038/s41396-021-00905-1

Zhu, X., Wang, J., Chen, Q., Chen, G., Huang, Y., & Yang, Z. (2016). Costs and trade-offs of grazer-induced defenses in Scenedesmus under deficient resource. Scientific Reports, 6(1), 22594. 10.1038/srep22594

