## Supplemental Tables for "Temperature stress disrupts reciprocal adaptation in a microbial predator-prey system"

| Population | Food Source | Doubling Time | Generations |
|---|---|---|---|
| ***T. pyriformis* Ancestor 26 °C** | Axenic Medium | 3.693 | 623.925 |
| ***T. pyriformis* Ancestor 30 °C** | Axenic Medium | 2.777 | 829.700 |
| ***T. pyriformis* Coevolved 26 °C** | Ancestral Bacteria | 9.620 | 239.504 |
| ***T. pyriformis* Coevolved 30 °C** | Ancestral Bacteria | 9.632 | 239.206 |
| ***T. pyriformis* Coevolved 26 °C** | Coevolved Bacteria | 9.927 | 232.084 |
| ***T. pyriformis* Coevolved 30 °C** | Coevolved Bacteria | 11.006 | 209.347 |
| ***P. fluorescens* Coevolved 30 °C** | Axenic Medium | 2.264 | 508.896 |
| ***P. fluorescens* Ancestor 26 °C** | Axenic Medium | 1.500 | 768.199 |
| ***P. fluorescens* Ancestor 30 °C** | Axenic Medium | 1.568 | 734.516 |

**Table S1:** Estimated number of generations that occurred during the evolution experiment under different conditions. Since there was little change in growth rates of evolved populations of either species, number of generations were estimated using ancestral generation times. Number of generations was calculated using the formula $\frac{Hours\ spent\ in\ exponetnial\ phas}{doubling\ time}$ for each condition. We estimated that bacterial and ciliate populations reached stationary phase by 24 or 48 hours respectively. We then estimated hours spent in exponential phase by multiplying the amount of time it took to reach stationary phase by the number of subcultures resulting in 1152 hours for *P. fluorescens* and 2304 hours for *T. pyriformis*. Doubling times for species grown on axenic medium come from growth rate measurements. We emphasize that these are rough estimates due to the uncertainties regarding extended periods of time in stationary phase between subcultures, differences in predator growth on axenic medium vs. bacteria, and population reduction in prey due to predation. Estimates were additionally obtained by calculating generations using growth rates for bacterial populations that did show changes in growth rate as well as for coevolved predator populations while fed bacteria.

| Population | Day | $\bar{D}$ | SE |
|---|---|---|---|
| **26 Alone 1** | 28 | -0.008 | 0.055 |
| **26 Alone 2** | 28 | 0.049 | 0.083 |
| **26 Alone 3** | 28 | 0.074 | 0.069 |
| **26 Alone 4** | 28 | 0.077 | 0.054 |
| **26 Alone 1** | 56 | 0.053 | 0.042 |
| **26 Alone 2** | 56 | -0.004 | 0.041 |
| **26 Alone 3** | 56 | 0.088 | 0.038 |
| **26 Alone 4** | 56 | -0.018 | 0.007 |
| **26 Alone 1** | 87 | 0.103 | 0.114 |
| **26 Alone 2** | 87 | 0.047 | 0.065 |
| **26 Alone 3** | 87 | 0.086 | 0.051 |
| **26 Alone 4** | 87 | 0.103 | 0.074 |
| **26 Alone 1** | 115 | -0.029 | 0.033 |
| **26 Alone 2** | 115 | -0.150 | 0.081 |
| **26 Alone 3** | 115 | -0.175 | 0.099 |
| **26 Alone 4** | 115 | 0.010 | 0.083 |
| **26 Alone 1** | 149 | -0.211 | 0.061 |
| **26 Alone 2** | 149 | -0.043 | 0.096 |
| **26 Alone 3** | 149 | -0.036 | 0.030 |
| **26 Alone 4** | 149 | -0.083 | 0.075 |
| **26 Alone 1** | 181 | -0.046 | 0.062 |
| **26 Alone 2** | 181 | 0.004 | 0.066 |
| **26 Alone 3** | 181 | -0.122 | 0.028 |
| **26 Alone 4** | 181 | 0.011 | 0.076 |
| **26 Together 1** | 28 | 0.348 | 0.009 |
| **26 Together 2** | 28 | 0.497 | 0.040 |
| **26 Together 3** | 28 | 0.422 | 0.024 |
| **26 Together 4** | 28 | 0.422 | 0.016 |
| **26 Together 1** | 56 | 0.492 | 0.025 |
| **26 Together 2** | 56 | 0.624 | 0.023 |
| **26 Together 3** | 56 | 0.573 | 0.017 |
| **26 Together 4** | 56 | 0.600 | 0.023 |
| **26 Together 1** | 87 | 0.419 | 0.033 |
| **26 Together 2** | 87 | 0.522 | 0.016 |
| **26 Together 3** | 87 | 0.541 | 0.028 |
| **26 Together 4** | 87 | 0.585 | 0.027 |
| **26 Together 1** | 115 | 0.645 | 0.009 |
| **26 Together 2** | 115 | 0.724 | 0.008 |
| **26 Together 3** | 115 | 0.757 | 0.028 |
| **26 Together 4** | 115 | 0.780 | 0.018 |
| **26 Together 1** | 149 | 0.644 | 0.013 |
| **26 Together 2** | 149 | 0.668 | 0.017 |
| **26 Together 3** | 149 | 0.634 | 0.015 |

| Population | Day | $\bar{D}$ | SE |
|---|---|---|---|
| **30 Alone 1** | 28 | 0.230 | 0.141 |
| **30 Alone 2** | 28 | 0.317 | 0.074 |
| **30 Alone 3** | 28 | 0.050 | 0.051 |
| **30 Alone 4** | 28 | -0.019 | 0.062 |
| **30 Alone 1** | 59 | -0.353 | 0.089 |
| **30 Alone 2** | 59 | -0.141 | 0.108 |
| **30 Alone 3** | 59 | -0.318 | 0.100 |
| **30 Alone 4** | 59 | -0.235 | 0.049 |
| **30 Alone 1** | 87 | -0.143 | 0.033 |
| **30 Alone 2** | 87 | -0.122 | 0.084 |
| **30 Alone 3** | 87 | -0.224 | 0.111 |
| **30 Alone 4** | 87 | 0.143 | 0.024 |
| **30 Alone 1** | 115 | -0.053 | 0.054 |
| **30 Alone 2** | 115 | -0.295 | 0.082 |
| **30 Alone 3** | 115 | -0.265 | 0.049 |
| **30 Alone 4** | 115 | -0.015 | 0.059 |
| **30 Alone 1** | 143 | 0.141 | 0.022 |
| **30 Alone 2** | 143 | 0.081 | 0.007 |
| **30 Alone 3** | 143 | -0.154 | 0.029 |
| **30 Alone 4** | 143 | -0.087 | 0.013 |
| **30 Together 1** | 171 | -0.127 | 0.056 |
| **30 Together 2** | 171 | -0.038 | 0.025 |
| **30 Together 3** | 171 | -0.068 | 0.073 |
| **30 Together 4** | 171 | 0.030 | 0.008 |
| **30 Together 1** | 28 | 0.205 | 0.050 |
| **30 Together 2** | 28 | 0.149 | 0.036 |
| **30 Together 3** | 28 | 0.317 | 0.039 |
| **30 Together 4** | 28 | 0.311 | 0.073 |
| **30 Together 1** | 59 | 0.434 | 0.026 |
| **30 Together 2** | 59 | 0.358 | 0.032 |
| **30 Together 3** | 59 | 0.302 | 0.012 |
| **30 Together 4** | 59 | 0.409 | 0.049 |
| **30 Together 1** | 87 | 0.372 | 0.023 |
| **30 Together 2** | 87 | 0.386 | 0.031 |
| **30 Together 3** | 87 | 0.489 | 0.046 |
| **30 Together 4** | 87 | 0.544 | 0.016 |
| **30 Together 1** | 115 | 0.373 | 0.016 |
| **30 Together 2** | 115 | 0.382 | 0.025 |
| **30 Together 3** | 115 | 0.373 | 0.040 |
| **30 Together 4** | 115 | 0.358 | 0.022 |
| **30 Together 1** | 143 | 0.448 | 0.043 |
| **30 Together 2** | 143 | 0.443 | 0.025 |
| **30 Together 3** | 143 | 0.431 | 0.011 |

| | | | | | | | |
|---|---|---|---|---|---|---|---|
| **26 Together 4** | 149 | 0.705 | 0.012 | **30 Together 4** | 143 | 0.443 | 0.022 |
| **26 Together 1** | 181 | 0.760 | 0.025 | **30 Together 1** | 171 | 0.416 | 0.004 |
| **26 Together 2** | 181 | 0.799 | 0.028 | **30 Together 2** | 171 | 0.386 | 0.021 |
| **26 Together 3** | 181 | 0.789 | 0.014 | **30 Together 3** | 171 | 0.342 | 0.028 |
| **26 Together 4** | 181 | 0.778 | 0.011 | **30 Together 4** | 171 | 0.410 | 0.007 |

**Table S2:** Bacterial defense of all *P. fluorescens* populations over time. Mean defense ($\bar{\boldsymbol{D}}$) values were calculated as the average of 4 technical replicates per population, as well as the standard error of the mean (SE). Means of biological replicates of each condition are plotted in Figure 2.

| **Population** | **Day** | $\overline{\boldsymbol{FE}}$ | **SE** |
|---|---|---|---|
| **26 Alone 1** | 181 | 0.080 | 0.059 |
| **26 Alone 2** | 181 | 0.097 | 0.090 |
| **26 Alone 3** | 181 | 0.142 | 0.081 |
| **26 Alone 4** | 181 | -0.097 | 0.023 |
| **26 Together 1** | 181 | 0.333 | 0.051 |
| **26 Together 2** | 181 | 0.633 | 0.131 |
| **26 Together 3** | 181 | 0.589 | 0.076 |
| **26 Together 4** | 181 | 0.444 | 0.056 |
| **30 Alone 1** | 171 | 0.123 | 0.040 |
| **30 Alone 2** | 171 | 0.144 | 0.110 |
| **30 Alone3** | 171 | 0.068 | 0.046 |
| **30 Alone 4** | 171 | 0.014 | 0.087 |
| **30 Together 1** | 171 | 0.000 | 0.039 |
| **30 Together 2** | 171 | 0.304 | 0.054 |
| **30 Together 3** | 171 | 0.348 | 0.040 |
| **30 Together 4** | 171 | 0.161 | 0.045 |

**Table S3:** Ciliate feeding efficiency of all *T. pyriformis* populations over time. Mean feeding efficiency values (**$\overline{\boldsymbol{FE}}$**) were calculated as the average of 4 technical replicates per population, as well as the standard error of the mean (SE). Means of biological replicates of each condition are plotted in Figure 3.

| Population | Day | $\bar{r}$ | SE |
|---|---|---|---|
| **26 Alone 1** | 28 | 0.425 | 0.007 |
| **26 Alone 2** | 28 | 0.410 | 0.008 |
| **26 Alone 3** | 28 | 0.394 | 0.006 |
| **26 Alone 4** | 28 | 0.386 | 0.005 |
| **26 Alone 1** | 87 | 0.415 | 0.007 |
| **26 Alone 2** | 87 | 0.438 | 0.002 |
| **26 Alone 3** | 87 | 0.427 | 0.004 |
| **26 Alone 4** | 87 | 0.412 | 0.008 |
| **26 Alone 1** | 115 | 0.402 | 0.003 |
| **26 Alone 2** | 115 | 0.405 | 0.004 |
| **26 Alone 3** | 115 | 0.431 | 0.003 |
| **26 Alone 4** | 115 | 0.400 | 0.008 |
| **26 Alone 1** | 149 | 0.397 | 0.003 |
| **26 Alone 2** | 149 | 0.409 | 0.003 |
| **26 Alone 3** | 149 | 0.408 | 0.005 |
| **26 Alone 4** | 149 | 0.407 | 0.003 |
| **26 Alone 1** | 181 | 0.399 | 0.005 |
| **26 Alone 2** | 181 | 0.394 | 0.003 |
| **26 Alone 3** | 181 | 0.358 | 0.004 |
| **26 Alone 4** | 181 | 0.349 | 0.007 |
| **26 Together 1** | 28 | 0.330 | 0.004 |
| **26 Together 2** | 28 | 0.387 | 0.013 |
| **26 Together 4** | 28 | 0.375 | 0.004 |
| **26 Together 1** | 87 | 0.369 | 0.006 |
| **26 Together 2** | 87 | 0.421 | 0.002 |
| **26 Together 3** | 87 | 0.405 | 0.005 |
| **26 Together 4** | 87 | 0.341 | 0.004 |
| **26 Together 1** | 115 | 0.366 | 0.005 |
| **26 Together 2** | 115 | 0.331 | 0.008 |
| **26 Together 3** | 115 | 0.385 | 0.010 |
| **26 Together 4** | 115 | 0.327 | 0.008 |
| **26 Together 1** | 149 | 0.361 | 0.004 |
| **26 Together 2** | 149 | 0.409 | 0.008 |
| **26 Together 3** | 149 | 0.367 | 0.004 |
| **26 Together 4** | 149 | 0.387 | 0.013 |
| **26 Together 1** | 181 | 0.386 | 0.009 |
| **26 Together 2** | 181 | 0.411 | 0.005 |
| **26 Together 3** | 181 | 0.369 | 0.008 |
| **26 Together 4** | 181 | 0.352 | 0.005 |
| **26 Ancestor** | 28 | 0.390 | 0.003 |
| **26 Ancestor** | 87 | 0.428 | 0.005 |
| **26 Ancestor** | 115 | 0.409 | 0.004 |

| Population | Day | $\bar{r}$ | SE |
|---|---|---|---|
| **30 Alone 1** | 28 | 0.462 | 0.003 |
| **30 Alone 2** | 28 | 0.453 | 0.008 |
| **30 Alone 3** | 28 | 0.413 | 0.003 |
| **30 Alone 4** | 28 | 0.369 | 0.090 |
| **30 Alone 1** | 59 | 0.467 | 0.005 |
| **30 Alone 2** | 59 | 0.465 | 0.007 |
| **30 Alone 3** | 59 | 0.471 | 0.009 |
| **30 Alone 4** | 59 | 0.473 | 0.011 |
| **30 Alone 1** | 87 | 0.421 | 0.016 |
| **30 Alone 2** | 87 | 0.402 | 0.016 |
| **30 Alone 3** | 87 | 0.434 | 0.013 |
| **30 Alone 4** | 87 | 0.452 | 0.015 |
| **30 Alone 1** | 115 | 0.415 | 0.004 |
| **30 Alone 2** | 115 | 0.398 | 0.003 |
| **30 Alone 4** | 115 | 0.409 | 0.006 |
| **30 Alone 1** | 143 | 0.412 | 0.004 |
| **30 Alone 2** | 143 | 0.390 | 0.005 |
| **30 Alone 3** | 143 | 0.399 | 0.005 |
| **30 Alone 4** | 143 | 0.406 | 0.003 |
| **30 Alone 1** | 171 | 0.499 | 0.012 |
| **30 Alone 2** | 171 | 0.459 | 0.005 |
| **30 Alone3** | 171 | 0.492 | 0.005 |
| **30 Alone 4** | 171 | 0.482 | 0.015 |
| **30 Together 1** | 28 | 0.517 | 0.006 |
| **30 Together 2** | 28 | 0.480 | 0.008 |
| **30 Together 3** | 28 | 0.485 | 0.004 |
| **30 Together 4** | 28 | 0.463 | 0.006 |
| **30 Together 1** | 59 | 0.411 | 0.013 |
| **30 Together 2** | 59 | 0.395 | 0.008 |
| **30 Together 3** | 59 | 0.438 | 0.013 |
| **30 Together 4** | 59 | 0.411 | 0.005 |
| **30 Together 1** | 87 | 0.414 | 0.006 |
| **30 Together 2** | 87 | 0.426 | 0.008 |
| **30 Together 3** | 87 | 0.402 | 0.009 |
| **30 Together 4** | 87 | 0.414 | 0.011 |
| **30 Together 1** | 115 | 0.383 | 0.004 |
| **30 Together 2** | 115 | 0.382 | 0.003 |
| **30 Together 3** | 115 | 0.409 | 0.001 |
| **30 Together 4** | 115 | 0.393 | 0.004 |
| **30 Together 1** | 143 | 0.374 | 0.005 |
| **30 Together 2** | 143 | 0.391 | 0.006 |
| **30 Together 3** | 143 | 0.323 | 0.010 |

| | | | |
|---|---|---|---|
| **26 Ancestor** | 149 | 0.401 | 0.004 |
| **26 Ancestor** | 181 | 0.414 | 0.007 |
| **30 Together 1** | 171 | 0.427 | 0.007 |
| **30 Together 2** | 171 | 0.289 | 0.020 |
| **30 Together 3** | 171 | 0.149 | 0.007 |
| **30 Together 4** | 171 | 0.360 | 0.015 |
| **30 Ancestor** | 28 | 0.446 | 0.005 |
| **30 Ancestor** | 59 | 0.487 | 0.009 |
| **30 Ancestor** | 87 | 0.479 | 0.013 |
| **30 Ancestor** | 115 | 0.399 | 0.004 |
| **30 Ancestor** | 143 | 0.396 | 0.005 |
| **30 Ancestor** | 171 | 0.455 | 0.013 |

**Table S4:** Growth rates (divisions/hr) on axenic medium of all *P. fluorescens* populations over time. Mean growth rates ($\bar{\boldsymbol{r}}$) were calculated as the average of 5 technical replicates per population, as well as the standard error of the mean (SE).

| Population | Day | $\overline{\Delta r}$ |
|---|---|---|
| **26 Alone 1** | 28 | 0.035 |
| **26 Alone 2** | 28 | 0.020 |
| **26 Alone 3** | 28 | 0.004 |
| **26 Alone 4** | 28 | -0.005 |
| **26 Alone 1** | 87 | -0.013 |
| **26 Alone 2** | 87 | 0.010 |
| **26 Alone 3** | 87 | -0.001 |
| **26 Alone 4** | 87 | -0.016 |
| **26 Alone 1** | 115 | -0.007 |
| **26 Alone 2** | 115 | -0.004 |
| **26 Alone 3** | 115 | 0.022 |
| **26 Alone 4** | 115 | -0.008 |
| **26 Alone 1** | 149 | -0.005 |
| **26 Alone 2** | 149 | 0.007 |
| **26 Alone 3** | 149 | 0.006 |
| **26 Alone 4** | 149 | 0.006 |
| **26 Alone 1** | 181 | -0.015 |
| **26 Alone 2** | 181 | -0.020 |
| **26 Alone 3** | 181 | -0.056 |
| **26 Alone 4** | 181 | -0.065 |
| **26 Together 1** | 28 | -0.060 |
| **26 Together 2** | 28 | -0.003 |
| **26 Together 4** | 28 | -0.015 |
| **26 Together 1** | 87 | -0.059 |
| **26 Together 2** | 87 | -0.007 |
| **26 Together 3** | 87 | -0.023 |
| **26 Together 4** | 87 | -0.087 |
| **26 Together 1** | 115 | -0.043 |
| **26 Together 2** | 115 | -0.078 |
| **26 Together 3** | 115 | -0.023 |
| **26 Together 4** | 115 | -0.082 |
| **26 Together 1** | 149 | -0.040 |
| **26 Together 2** | 149 | 0.008 |
| **26 Together 3** | 149 | -0.035 |
| **26 Together 4** | 149 | -0.015 |
| **26 Together 1** | 181 | -0.028 |
| **26 Together 2** | 181 | -0.003 |
| **26 Together 3** | 181 | -0.045 |
| **26 Together 4** | 181 | -0.062 |

| Population | Day | $\overline{\Delta r}$ |
|---|---|---|
| **30 Alone 1** | 28 | 0.016 |
| **30 Alone 2** | 28 | 0.007 |
| **30 Alone 3** | 28 | -0.034 |
| **30 Alone 4** | 28 | -0.077 |
| **30 Alone 1** | 59 | -0.020 |
| **30 Alone 2** | 59 | -0.022 |
| **30 Alone 3** | 59 | -0.016 |
| **30 Alone 4** | 59 | -0.014 |
| **30 Alone 1** | 87 | -0.058 |
| **30 Alone 2** | 87 | -0.077 |
| **30 Alone 3** | 87 | -0.045 |
| **30 Alone 4** | 87 | -0.027 |
| **30 Alone 1** | 115 | 0.016 |
| **30 Alone 2** | 115 | -0.002 |
| **30 Alone 4** | 115 | 0.010 |
| **30 Alone 1** | 143 | 0.015 |
| **30 Alone 2** | 143 | -0.006 |
| **30 Alone 3** | 143 | 0.003 |
| **30 Alone 4** | 143 | 0.010 |
| **30 Alone 1** | 171 | 0.045 |
| **30 Alone 2** | 171 | 0.005 |
| **30 Alone3** | 171 | 0.037 |
| **30 Alone 4** | 171 | 0.027 |
| **30 Together 1** | 28 | 0.071 |
| **30 Together 2** | 28 | 0.033 |
| **30 Together 3** | 28 | 0.039 |
| **30 Together 4** | 28 | 0.016 |
| **30 Together 1** | 59 | -0.076 |
| **30 Together 2** | 59 | -0.091 |
| **30 Together 3** | 59 | -0.049 |
| **30 Together 4** | 59 | -0.076 |
| **30 Together 1** | 87 | -0.065 |
| **30 Together 2** | 87 | -0.053 |
| **30 Together 3** | 87 | -0.077 |
| **30 Together 4** | 87 | -0.065 |
| **30 Together 1** | 115 | -0.016 |
| **30 Together 2** | 115 | -0.017 |
| **30 Together 3** | 115 | 0.010 |
| **30 Together 4** | 115 | -0.007 |
| **30 Together 1** | 143 | -0.022 |
| **30 Together 2** | 143 | -0.006 |
| **30 Together 3** | 143 | -0.073 |

| | | |
|---|---|---|
| **30 Together 1** | 171 | -0.028 |
| **30 Together 2** | 171 | -0.165 |
| **30 Together 3** | 171 | -0.306 |
| **30 Together 4** | 171 | -0.095 |

**Table S5:** Change in growth rates (divisions/hr) on axenic medium of all *P. fluorescens* populations over time. Mean change in growth values ($\overline{\Delta r}$) were calculated as $\bar{r}_{Evolved} - \bar{r}_{Ancestor}$ at a given timepoint. Means of biological replicates of each condition are plotted in Figure 4.

| Population | Day | $\bar{r}$ | SE |
|---|---|---|---|
| **26 Alone 1** | 28 | 0.210 | 0.004 |
| **26 Alone 2** | 28 | 0.211 | 0.004 |
| **26 Alone 3** | 28 | 0.215 | 0.003 |
| **26 Alone 4** | 28 | 0.219 | 0.004 |
| **26 Alone 1** | 56 | 0.169 | 0.003 |
| **26 Alone 2** | 56 | 0.139 | 0.003 |
| **26 Alone 3** | 56 | 0.126 | 0.029 |
| **26 Alone 4** | 56 | 0.138 | 0.006 |
| **26 Alone 1** | 181 | 0.205 | 0.006 |
| **26 Alone 2** | 181 | 0.196 | 0.005 |
| **26 Alone 3** | 181 | 0.203 | 0.001 |
| **26 Alone 4** | 181 | 0.195 | 0.006 |
| **26 Together 1** | 28 | 0.204 | 0.007 |
| **26 Together 2** | 28 | 0.218 | 0.004 |
| **26 Together 3** | 28 | 0.210 | 0.005 |
| **26 Together 4** | 28 | 0.202 | 0.005 |
| **26 Together 1** | 56 | 0.126 | 0.006 |
| **26 Together 2** | 56 | 0.136 | 0.007 |
| **26 Together 3** | 56 | 0.122 | 0.007 |
| **26 Together 4** | 56 | 0.155 | 0.007 |
| **26 Together 1** | 181 | 0.196 | 0.010 |
| **26 Together 2** | 181 | 0.123 | 0.005 |
| **26 Together 3** | 181 | 0.165 | 0.014 |
| **26 Together 4** | 181 | 0.200 | 0.002 |

| Population | Day | $\bar{r}$ | SE |
|---|---|---|---|
| **30 Alone 1** | 28 | 0.258 | 0.007 |
| **30 Alone 2** | 28 | 0.259 | 0.006 |
| **30 Alone 3** | 28 | 0.256 | 0.006 |
| **30 Alone 4** | 28 | 0.245 | 0.010 |
| **30 Alone 1** | 87 | 0.194 | 0.004 |
| **30 Alone 2** | 87 | 0.242 | 0.001 |
| **30 Alone 3** | 87 | 0.281 | 0.002 |
| **30 Alone 4** | 87 | 0.249 | 0.003 |
| **30 Alone 1** | 171 | 0.231 | 0.005 |
| **30 Alone 2** | 171 | 0.225 | 0.004 |
| **30 Alone 3** | 171 | 0.229 | 0.003 |
| **30 Alone 4** | 171 | 0.230 | 0.004 |
| **30 Together 1** | 28 | 0.250 | 0.009 |
| **30 Together 2** | 28 | 0.253 | 0.010 |
| **30 Together 3** | 28 | 0.249 | 0.009 |
| **30 Together 4** | 28 | 0.256 | 0.004 |
| **30 Together 1** | 87 | 0.214 | 0.008 |
| **30 Together 2** | 87 | 0.252 | 0.003 |
| **30 Together 3** | 87 | 0.264 | 0.005 |
| **30 Together 4** | 87 | 0.288 | 0.007 |
| **30 Together 1** | 171 | 0.210 | 0.003 |
| **30 Together 2** | 171 | 0.235 | 0.014 |
| **30 Together 3** | 171 | 0.203 | 0.007 |
| **30 Together 4** | 171 | 0.171 | 0.011 |

**Table S6:** Growth rates (divisions/hr) on axenic medium of all *T. pyriformis* populations over time. Mean growth rates ($\bar{r}$) were calculated as the average of 5 technical replicates per population, as well as the standard error of the mean (SE).

| Population | Day | $\overline{\Delta r}$ |
|---|---|---|
| **26 Alone 1** | 28 | -0.002 |
| **26 Alone 2** | 28 | -0.001 |
| **26 Alone 3** | 28 | 0.003 |
| **26 Alone 4** | 28 | 0.007 |
| **26 Alone 1** | 56 | 0.008 |
| **26 Alone 2** | 56 | -0.022 |
| **26 Alone 3** | 56 | -0.035 |
| **26 Alone 4** | 56 | -0.023 |
| **26 Alone 1** | 181 | 0.014 |
| **26 Alone 2** | 181 | 0.006 |
| **26 Alone 3** | 181 | 0.013 |
| **26 Alone 4** | 181 | 0.004 |
| **26 Together 1** | 28 | -0.008 |
| **26 Together 2** | 28 | 0.006 |
| **26 Together 3** | 28 | -0.001 |
| **26 Together 4** | 28 | -0.010 |
| **26 Together 1** | 56 | -0.035 |
| **26 Together 2** | 56 | -0.025 |
| **26 Together 3** | 56 | -0.039 |
| **26 Together 4** | 56 | -0.006 |
| **26 Together 1** | 181 | 0.005 |
| **26 Together 2** | 181 | -0.067 |
| **26 Together 3** | 181 | -0.025 |
| **26 Together 4** | 181 | 0.010 |

| Population | Day | $\overline{\Delta r}$ |
|---|---|---|
| **30 Alone 1** | 28 | -0.001 |
| **30 Alone 2** | 28 | 0.001 |
| **30 Alone 3** | 28 | -0.003 |
| **30 Alone 4** | 28 | -0.013 |
| **30 Alone 1** | 87 | -0.071 |
| **30 Alone 2** | 87 | -0.023 |
| **30 Alone 3** | 87 | 0.016 |
| **30 Alone 4** | 87 | -0.016 |
| **30 Alone 1** | 171 | 0.006 |
| **30 Alone 2** | 171 | -0.001 |
| **30 Alone 3** | 171 | 0.004 |
| **30 Alone 4** | 171 | 0.005 |
| **30 Together 1** | 28 | -0.008 |
| **30 Together 2** | 28 | -0.005 |
| **30 Together 3** | 28 | -0.009 |
| **30 Together 4** | 28 | -0.002 |
| **30 Together 1** | 87 | -0.050 |
| **30 Together 2** | 87 | -0.013 |
| **30 Together 3** | 87 | -0.001 |
| **30 Together 4** | 87 | 0.023 |
| **30 Together 1** | 171 | -0.016 |
| **30 Together 2** | 171 | 0.010 |
| **30 Together 3** | 171 | -0.023 |
| **30 Together 4** | 171 | -0.055 |

**Table S7:** Change in growth rates (divisions/hr) on axenic medium of all *T. pyriformis* populations over time. Mean change in growth values ($\overline{\Delta r}$) were calculated as $\bar{r}_{Evolved} - \bar{r}_{Ancestor}$ at a given timepoint. Means of biological replicates of each condition are plotted in Figure 5A.

| Population | Bacteria | $\overline{DT}$ | SE |
|---|---|---|---|
| **26 Together 1** | Ancestral | 9.935 | 0.114 |
| **26 Together 2** | Ancestral | 9.395 | 0.200 |
| **26 Together 3** | Ancestral | 9.446 | 0.134 |
| **26 Together 4** | Ancestral | 9.703 | 0.106 |
| **26 Ancestor** | Ancestral | 10.519 | 0.198 |
| **26 Together 1** | 26 Coevolved 1 | 10.344 | 0.119 |
| **26 Together 2** | 26 Coevolved 2 | 10.047 | 0.195 |
| **26 Together 3** | 26 Coevolved 3 | 9.365 | 0.145 |
| **26 Together 4** | 26 Coevolved 4 | 9.954 | 0.207 |
| **26 Ancestor** | 26 Coevolved 1 | 10.913 | 0.374 |
| **26 Ancestor** | 26 Coevolved 2 | 11.672 | 0.526 |
| **26 Ancestor** | 26 Coevolved 3 | 11.336 | 0.267 |
| **26 Ancestor** | 26 Coevolved 4 | 11.130 | 0.196 |
| **30 Together 1** | Ancestral | 10.145 | 0.124 |
| **30 Together 2** | Ancestral | 9.387 | 0.110 |
| **30 Together 3** | Ancestral | 9.293 | 0.077 |
| **30 Together 4** | Ancestral | 9.703 | 0.106 |
| **30 Ancestor** | Ancestral | 9.664 | 0.168 |
| **30 Together 1** | 30 Coevolved 1 | 10.875 | 0.195 |
| **30 Together 2** | 30 Coevolved 2 | 11.548 | 0.398 |
| **30 Together 3** | 30 Coevolved 3 | 10.553 | 0.246 |
| **30 Together 4** | 30 Coevolved 4 | 11.047 | 0.329 |
| **30 Ancestor** | 30 Coevolved 1 | 10.306 | 0.025 |
| **30 Ancestor** | 30 Coevolved 2 | 10.156 | 0.106 |
| **30 Ancestor** | 30 Coevolved 3 | 9.950 | 0.129 |
| **30 Ancestor** | 30 Coevolved 4 | 10.277 | 0.038 |

**Table S8:** Doubling times (hrs) of coevolved *T. pyriformis* using bacterial prey. Mean doubling times ($\overline{DT}$) were calculated as the average of 4 technical replicates per population, as well as the standard error of the me (SE).

| Population | Bacteria | $\overline{\Delta DT}$ |
|---|---|---|
| **26 Together 1** | Ancestral | -0.584 |
| **26 Together 2** | Ancestral | -1.124 |
| **26 Together 3** | Ancestral | -1.072 |
| **26 Together 4** | Ancestral | -0.816 |
| **26 Together 1** | 26 Coevolved 1 | -0.569 |
| **26 Together 2** | 26 Coevolved 2 | -1.625 |
| **26 Together 3** | 26 Coevolved 3 | -1.971 |
| **26 Together 4** | 26 Coevolved 4 | -1.177 |
| **30 Together 1** | Ancestral | 0.481 |
| **30 Together 2** | Ancestral | -0.277 |
| **30 Together 3** | Ancestral | -0.371 |
| **30 Together 4** | Ancestral | 0.039 |
| **30 Together 1** | 26 Coevolved 1 | 0.569 |
| **30 Together 2** | 26 Coevolved 2 | 1.392 |
| **30 Together 3** | 26 Coevolved 3 | 0.603 |
| **30 Together 4** | 26 Coevolved 4 | 0.770 |

**Table S9:** Change in doubling times (hrs) of coevolved *T. pyriformis* using bacterial prey. Mean change in growth values ( $\overline{\Delta DT}$ ) were calculated as $\overline{\Delta DT}_{Evolved} - \overline{\Delta DT}_{Ancestor}$ for each bacterial prey type. Means of biological replicates of each condition are plotted in Figure 5B.
